# CRISPR/Cas9 screen for genome-wide interrogation of essential MYC binding sites in cancer cells

**DOI:** 10.1101/2021.08.02.454734

**Authors:** Marta Kazimierska, Marta Podralska, Magdalena Żurawek, Tomasz Woźniak, Marta Elżbieta Kasprzyk, Weronika Sura, Wojciech Łosiewski, Iwona Ziółkowska-Suchanek, Joost Kluiver, Anke van den Berg, Natalia Rozwadowska, Agnieszka Dzikiewicz-Krawczyk

## Abstract

The transcription factor MYC is a proto-oncogene with a well-documented essential role in the pathogenesis and maintenance of several types of cancer. MYC binds to specific E-box sequences in the genome to regulate gene expression in a cell type- and developmental stage-specific manner. To date, a comprehensive analysis of direct MYC targets with essential roles in different types of cancer is missing. To enable identification of functional MYC binding sites and corresponding target genes, we designed a CRISPR/Cas9 library to destroy E-box sequences in a genome-wide fashion. In parallel, we used the Brunello library to knockout protein-coding genes. We performed high-throughput screens with these libraries in four MYC-dependent cancer cell lines: K562, ST486, HepG2 and MCF7, which revealed several essential E-boxes and genes. Among them we pinpointed crucial known and novel MYC-regulated genes involved in pathways associated with cancer development. Extensive validation of our approach in K562 cells confirmed that E-box disruption affects MYC binding, target genes expression and cell proliferation. Our unique, well-validated tool opens new possibilities to gain novel insights into MYC-dependent vulnerabilities in cancer cells.

## INTRODUCTION

MYC is an oncogene broadly involved in many tumors. Due to amplifications, mutations, translocations or posttranslational modifications, MYC is highly expressed in up to 70% of cancers^1^. The family of MYC transcription factors (TFs) consists of three members: C-MYC, N-MYC, and L-MYC^2^. Among them, C-MYC (further called MYC) has the strongest oncogenic potential and is widely deregulated in cancer, while N-MYC and L-MYC are mainly involved in neuroblastoma and lung cancer, respectively^3–5^. MYC regulates expression of genes associated with cell cycle, apoptosis, proliferation, cellular differentiation as well as strongly alters metabolism of cancer cells and stimulates ribosome and mitochondrial biogenesis^6–9^. Altogether, this creates strong addiction of cancer cells to MYC and it has been demonstrated that indeed MYC withdrawal leads to tumor regression^10^. Thus, targeting MYC appears as an attractive strategy for cancer therapies. However, no clinically relevant MYC-targeting therapies have been developed so far^11^. MYC is considered an “undruggable target” due to its localization and activity in the nucleus and lack of an active site for interaction with small molecules^2,4^. Therefore, there is a need to look for indirect approaches such as identification of essential MYC-regulated genes which may serve as a basis for novel anti-tumor therapies.

MYC is a TF belonging to the basic helix-loop-helix-leucine-zipper family (bHLH-LZ) that creates heterodimers with the MYC-associated factor X (MAX). The MYC/MAX complex recognizes and binds to E-box motifs in DNA (canonical sequence 5’-CACGTG-3’), localized mainly in promoters and enhancers. Although MYC has been widely studied, its targetome is not fully known. Thousands of genes responding to MYC activation or inhibition have been identified but it is difficult to distinguish direct and indirect targets^12–15^. Chromatin immunoprecipitation with MYC antibody followed by sequencing (MYC-ChIP-seq) indicated thousands of MYC-bound sites in the genome^12,16–19^. However, the set of MYC target genes varies among different cell types and developmental stages^20,21^. This may be explained by the fact that MYC binds predominantly to already active promoters or enhancers and inactive genes remain silent^16^. While it was shown that disruption of even one gene, crucial for MYC-dependent cancer development can be sufficient to decrease cancer cell growth^22^, it is still not clear which MYC targets are essential for cancer cells. To date, a limited number of RNA interference screens for genes essential in MYC-driven tumors have been performed. However, they either focused only on a limited set of genes or were performed in normal cells with forced MYC overexpression^23–26^. This precludes their direct relevance for cancer cells. So far, no global, comprehensive analysis of MYC target genes essential for MYC-dependent cancer cells has been performed.

In this study, we created a tool for a comprehensive analysis of essential MYC binding sites and target genes in MYC-dependent cancer cells. We designed a sgRNA library for a genome-wide disruption of MYC binding sites and conducted a high-throughput screen in four MYC-dependent cell lines: K562 (chronic myelogenous leukemia, CML), ST486 (Burkitt lymphoma, BL), HepG2 (hepatocellular carcinoma) and MCF7 (breast cancer). In parallel, we utilized the Brunello library for genome-wide knockout of protein-coding genes. Overlapping data from both screens identified several known and novel MYC targets critical for those cells. Altogether, we established a unique, well-validated tool to identify MYC-regulated target genes relevant for growth of malignant cells. Our findings provide novel insights into MYC-dependent vulnerabilities in cancer cells.

## RESULTS

### Generation of the MYC-CRISPR library for a genome-wide disruption of MYC binding sites

From publicly available MYC-ChIP-seq data in MYC-dependent K562, MCF7, HepG2^27^, and BL cell lines^18^ we obtained 58 503 MYC binding sites, which contained 43 153 E-box motifs. 2 208 of them were located in the coding exons and were excluded to prevent effects due to disruption of the protein. Based on the presence of PAM sequence, we designed 56 688 unique sgRNAs targeting the remaining 26 653 E-boxes (65.1%). After excluding sgRNAs with predicted off-target binding, we finally obtained 45 350 sgRNAs targeting 24 981 E-boxes (61%) (Figure 1A, Supplementary Table S1). Half of the E-boxes were targeted by more than one sgRNA (Figure 1B). The majority of E-boxes were located in introns (58.4%) or intergenic regions (33.2%) (Figure 1C). 61% of the E-boxes targeted by our MYC-CRISPR library were bound by MYC only in one cell line, while 6% of the E-boxes were commonly bound by MYC in all four cell lines (Figure 1D). This is in line with the high cell-type specificity of MYC targets. Although the MYC-CRISPR library was designed based on the data from selected four cell lines, analysis of available MYC-ChIP-seq data from 11 cell lines revealed that 16-60% MYC binding sites were targeted by the library sgRNAs (Figure 1E). This indicates that the MYC-CRISPR library can be widely applied for studies in various cell lines.

**Figure 1.**
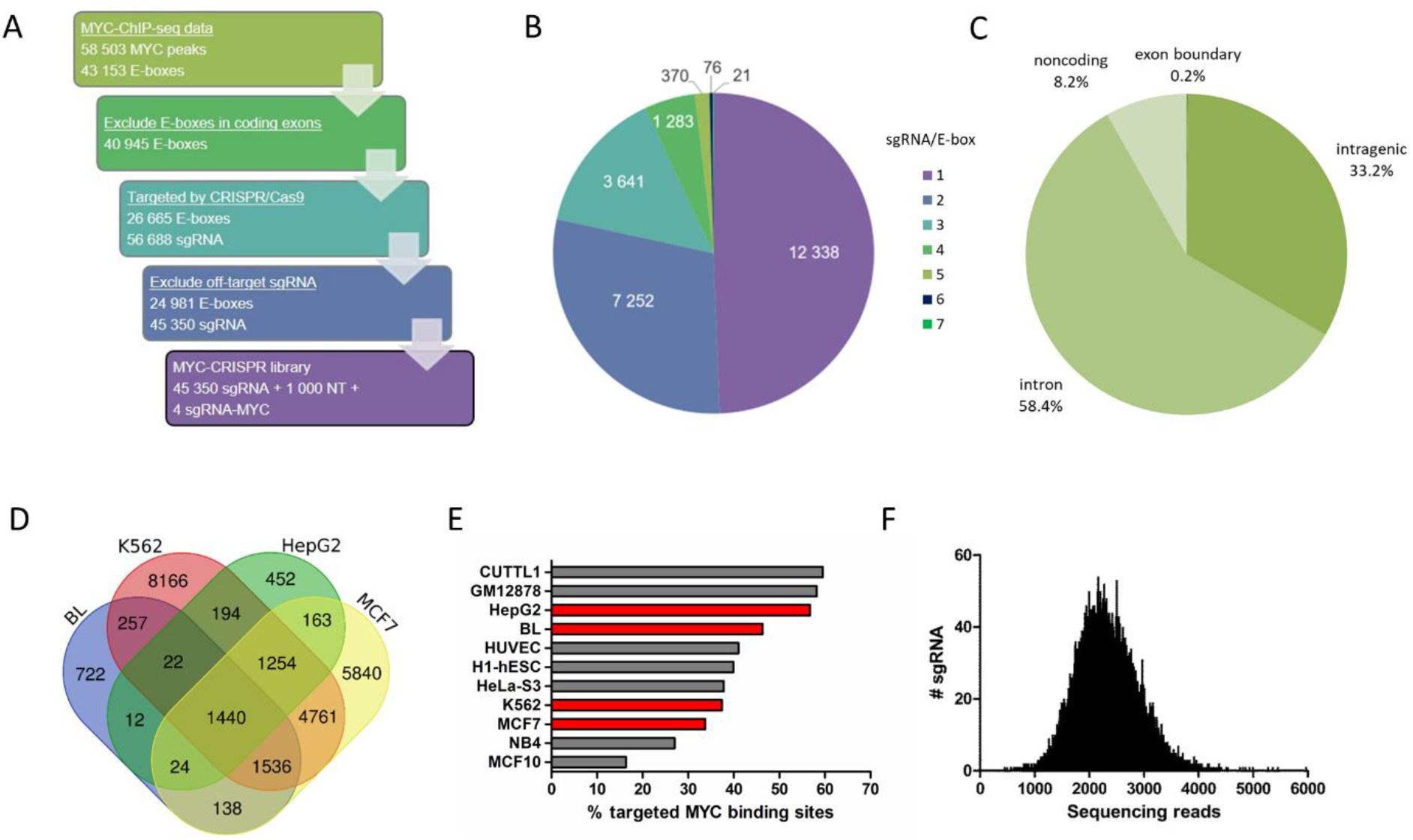
Design and generation of the MYC-CRISPR library for genome-wide disruption of MYC binding sites. **A**) Library was designed based on publicly available MYC-ChIP-seq data in MYC-dependent K562, MCF7, HepG2 and Burkitt lymphoma (BL) cell lines. After excluding E-boxes in coding exons, all possible sgRNAs targeting remaining E-boxes were designed, based on the presence of PAM sequence. sgRNAs with predicted off-target binding were filtered out. Final library contains 43,350 sgRNAs targeting E-boxes, 1,000 non-targeting (NT) sgRNAs as a negative control, and four sgRNAs targeting MYC as a positive control. **B**) Number of sgRNA constructs per E-box. **C**) Genomic location of E-boxes targeted by the library. **D**) Overlap of targeted E-boxes for selected cancer cell lines. **E**) Percentage of MYC binding sites targeted by the MYC-CRISPR library in various cell lines, based on available MYC-ChIP-Seq data. In red are cell lines for which the library was designed. **F**) Distribution of sgRNA constructs in the MYC-CRISPR plasmid library determined by NGS. All sgRNAs were present in the library.

Next generation sequencing revealed high quality of the cloned MYC-CRISPR library. All sgRNAs were present and the skew ratio of 90^th^ to 10^th^ percentile was only 1.76, indicating a uniform representation of all constructs in the library (Figure 1F). The quality of the amplified Brunello library also conformed to the recommended requirements (Supplementary Figure S1A-S1C). Thus, we generated a high quality MYC-CRISPR library for genome-wide targeting of MYC binding sites that can be universally used in various cell lines.

### High-throughput screen for functional MYC binding sites and target genes essential for growth of cancer cells

To identify essential MYC binding sites and target genes, we conducted a genome-wide high-throughput CRISPR/Cas9 screen with the MYC-CRISPR library (46 354 sgRNA) targeting E-boxes and the Brunello library (77 441 sgRNA) ^28^ targeting protein-coding genes (Figure 2A). Four cancer cell lines were infected in duplicate aiming at ~500x coverage of each sgRNA in both libraries and a 30% transduction efficiency. The obtained values are shown in Table 1. Cells were collected at T0 (after puromycin selection) and T1 (after 20 population doublings) and the abundance of sgRNA constructs was determined by NGS (Figure 2B, Supplementary Table S2, S3). Almost none of the non-targeting sgRNAs was depleted ≥2-fold in both replicates and 2-4 sgRNAs targeting MYC showed consistently decreased abundance at T1. 517-3 034 sgRNAs from the MYC-CRISPR library and 9 112-11 970 sgRNAs from the Brunello library were consistently ≥2-fold depleted in both replicates (Supplementary Figure 2A, 2B). Considering the combined effect of all sgRNAs targeting a given gene or E-box, using DESeq2 algorithm, 354-1 992 genes (Supplementary Table S4) and 56-97 E-boxes (Supplementary Table S5) were identified as essential for growth of selected cancer cells (p_adj_<0.001), while 3-9 E-boxes and 5-18 genes were significantly enriched (Figure 2C, 2D).

**Figure 2.**
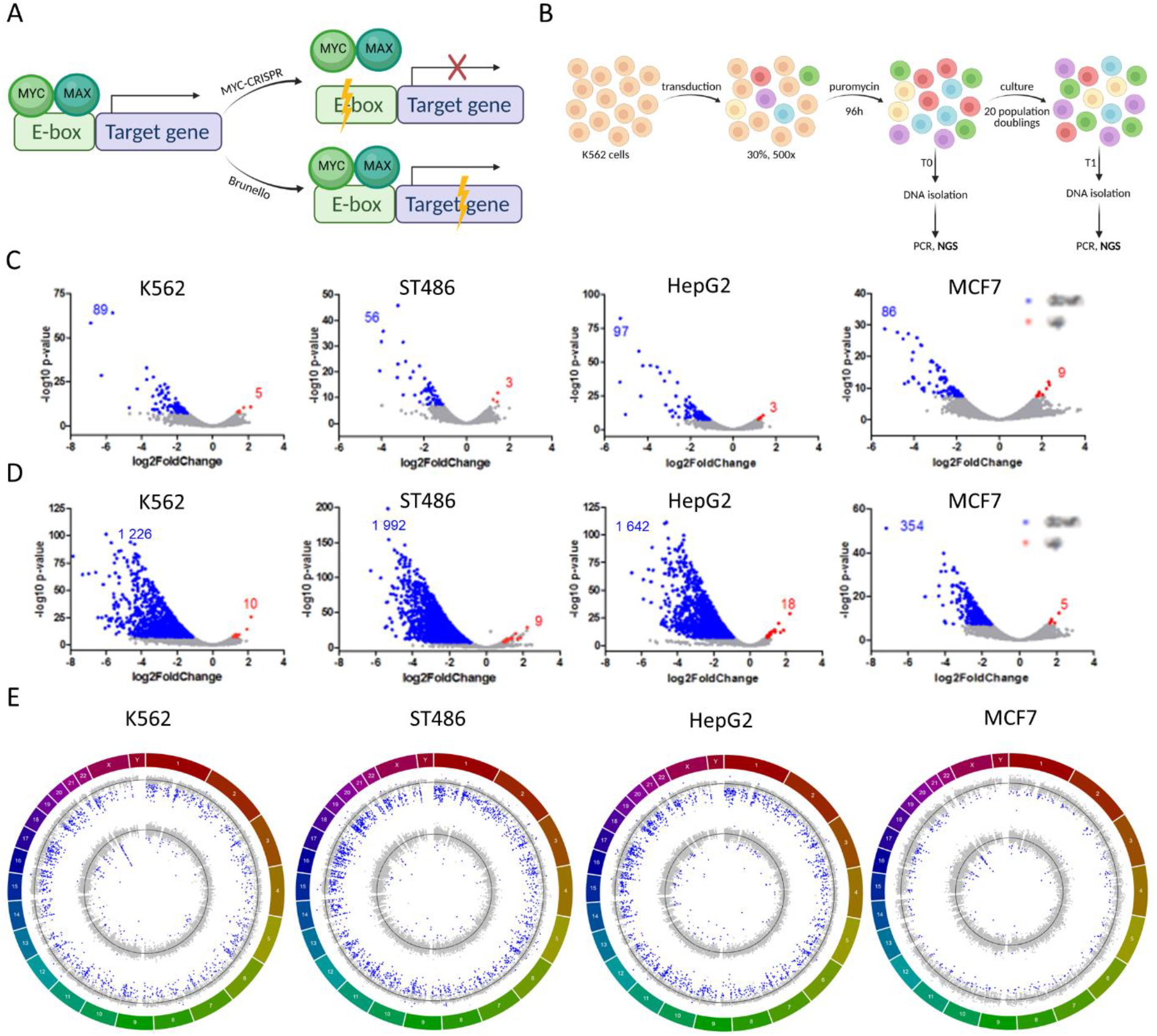
High-throughput screen with MYC-CRISPR and Brunello libraries. **A**) Experimental approach: 1) MYC-CRISPR library to destroy E-box sequences, disrupt MYC binding and its effect on target gene expression 2) Brunello library for genome-wide gene knock-out. **B**) Scheme of the high-throughput screen in cancer cells with MYC-CRISPR and Brunello libraries. **C**) DESeq2 analysis revealed essential E-boxes in MYC-CRISPR and **D**) essential genes in Brunello library (depleted genes in blue, enriched genes in red). **E**) Circos plots showing log2FC values for genes in Brunello screen (outer circle) and E-boxes in MYC-CRISPR screen (inner circle) across the chromosomes. Blue dots indicate genes and E-boxes significantly (p_adj_<0.001) depleted or enriched, black lines denote log2FC=0.

**Table 1.** Performance of the screen in MYC-dependent cancer cell lines conducted in duplicate.

|  | K562 |  | ST486 |  | HepG2 |  | MCF7 |  |
| --- | --- | --- | --- | --- | --- | --- | --- | --- |
|  | Transduction efficiency | Coverage | Transduction efficiency | Coverage | Transduction efficiency | Coverage | Transduction efficiency | Coverage |
| **MYC-CRISPR #1** | 32% | 540x | 26.2% | 435x | 29% | 485x | 30% | 500x |
| **MYC-CRISPR #2** | 26.9% | 455x | 32.1% | 535x | 33.3% | 560x | 25.4% | 425x |
| **Brunello #1** | 23% | 390x | 37.5% | 625x | 26.2% | 440x | 24.1% | 405x |
| **Brunello #2** | 24.5% | 415x | 34.6% | 575x | 24.1% | 405x | 23.7% | 400x |

Initial analysis in K562 cell line revealed 152 essential E-boxes. 40% of them were localized on chromosome 22, near the breakpoint in *BCR* involved in the t(9;22) translocation (Figure 2E). Hits observed in this region are most likely not caused by targeting essential genes but due to massive CRISPR/Cas9-mediated DNA cleavage within this tandemly amplified region in K562 cells, as observed previously^29^. Indeed, an orthogonal approach with dCas9 which does not induce DNA cleavage but blocks E-box sites to prevent MYC binding demonstrated no effect on K562 cell growth for two sgRNAs from chromosome 22q11 (Supplementary Figure S3A). Therefore E-boxes from the amplified region on 22q11 and 9q34 were excluded from further analysis.

Analysis of essential E-boxes revealed that 20-32% were localized close to genes essential for cancer cells. Moreover, 42-49% of adjacent genes are well known MYC-regulated targets (Supplementary Table S5).

Thus, our CRISPR/Cas9 screen revealed known and novel MYC-dependent vulnerabilities in cancer cells.

### Common and cell type-specific MYC-regulated processes

Overlap of essential E-boxes for four cancer cell lines revealed only three (1%) common E-boxes (Supplementary Figure S4A): chr1_BS1363_CACAATG with neighbor genes *MECR* and *PTPRU*, chr11_BS79_CGCGTG localized near *RPLP2* and *PIDD1*, and chr18_BS691_CATGTG adjacent to *RBFA* and *TXNL4A*. 10 E-boxes (3%) overlapped in 3 out of 4 cell lines, while majority of E-boxes (59-87%) were essential only in one cell line. Crucial genes identified in the Brunello screen showed greater overlap, with 135 (5%) genes common for all cell lines and 788 (29%) in 3 out of 4, and only 10-30% cell type-specific genes (Supplementary Figure S4B).

To determine main functions of essential and MYC-regulated genes identified in our high-throughput screens we conducted Gene Ontology (GO) and Gene Set Enrichment (GSEA) analyses. Genes essential in the Brunello screen were involved in several GO processes common for all cell lines, such as: metabolism, ribosome biogenesis, metabolism of nucleic acids, splicing and translation (Figure 3A). GSEA results revealed very similar processes, majority of which were shared between cell lines. In addition, some cell-specific processes emerged, such as DNA repair in K562, aminoacyl tRNA biosynthesis in ST486, oxidative phosphorylation in HepG2 and cell cycle in MCF7 (Figure 3B, Supplementary Table S6). On the other hand, genes localized near depleted E-boxes showed a more diverse spectrum of processes, reflecting the limited overlap from the screen and indicating different processes regulated by MYC. We identified GO processes such as metabolism and ribosome biogenesis but also histone modifications, protein localization, RNA processing and metabolism (Figure 3C). GSEA for genes nearby E-boxes highlighted translation, ribosome biogenesis, RNA processing and tumor invasiveness as processes common for all cell lines. Cell type-specific processes were much more prevalent and diverse, with no particular predominant terms emerging (Figure 3D, Supplementary Table S7). Noteworthy, both in Brunello and MYC-CRISPR analyses, the HALLMARK_MYC_TARGETS was among the top enriched gene sets, which confirms accuracy of our approach. Interestingly, REACTOME_REGULATION_OF_EXPRESSION_OF_SLITS_AND_ROBOS was a recurrently enriched gene set in all cell lines, in Brunello as well as MYC-CRISPR results. SLIT/ROBO pathway is involved in axon guidance and cell migration but it has been also implicated in tumor growth, migration, angiogenesis, and microenvironment^30^.

**Figure 3.**
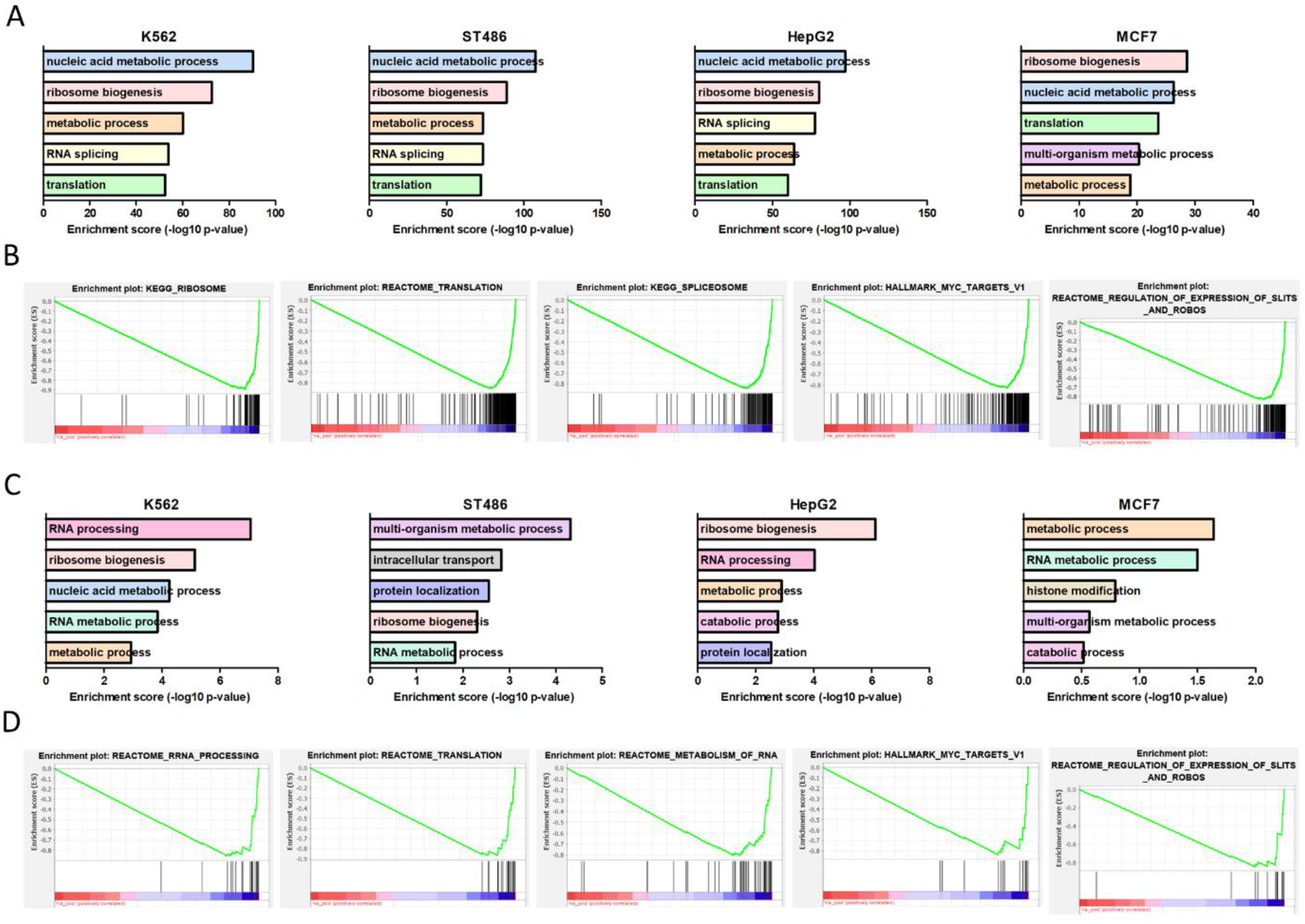
Essential MYC-regulated processes and pathways. A) GO analysis for essential genes from Brunello library. B) Selected GSEA gene sets enriched in Brunello screen in all cancer cell lines, example plots from K562 cell line are shown. C) GO analysis for genes localized up to 50 kb from essential E-boxes. D) Selected GSEA gene sets enriched in MYC-CRISPR screen in all cancer cell lines, example plots from K562 cell line are shown.

### Validation of the approach

To validate the results of the screen and confirm robustness of our approach for identification of MYC-dependent vulnerabilities in cancer cells, we focused on the top 10 most significantly depleted E-boxes in K562 cells. Of these, we included for validation six that were located nearby a protein-coding gene that was at least 4-fold depleted in Brunello screen (chr3_BS897_CATGTG, chr11_BS79_CGCGTG, chr11_BS2113_CACATG, chr13_BS121_CGCGTG, chr17_BS377_CACGTG, chr19_BS2255_CACATG), and two E-boxes adjacent to long non-coding RNA (lncRNA) genes (chr10_BS212_CACATG, chr2_BS1664_CACGTG). The remaining two of top 10 E-boxes were in vicinity of non-essential genes and were not considered for validation. In addition, we also included for validation a non-essential E-box, chr17_BS377_CGCGTG which was located 12 nt downstream of chr17_BS377_CACGTG (Supplementary Figure S5C), to gain further insight into MYC regulation at this locus.

Cells were transduced with individual sgRNAs targeting selected E-boxes. First, we checked whether our approach allows for efficient disruption of E-box motifs. TIDE analysis confirmed DNA editing with >90% efficiency for all sgRNAs. The spectrum of mutations varied between individual constructs, with small 1-2 nt indels being most prevalent (Figure 4A).

**Figure 4.**
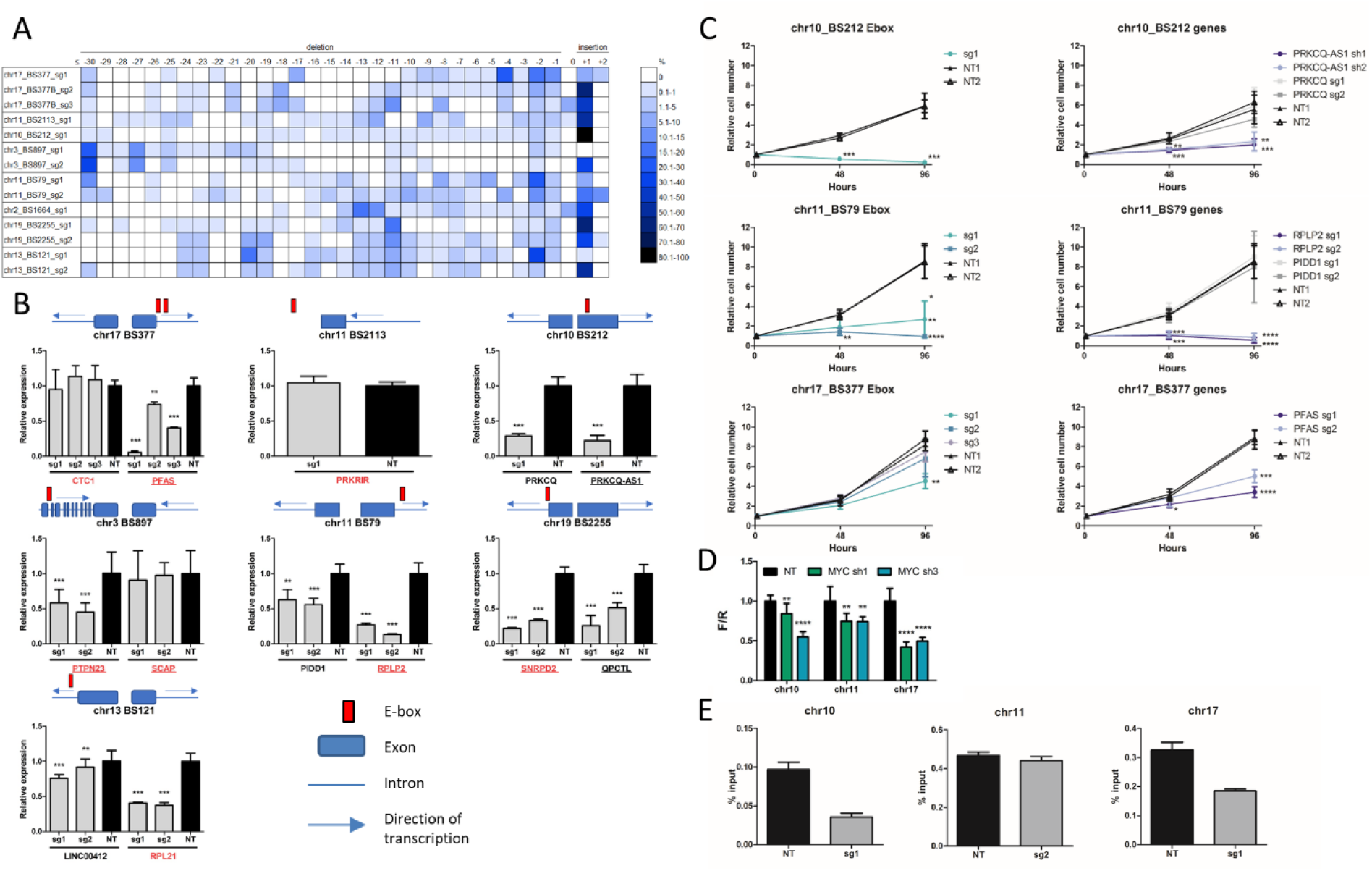
Validation of selected E-boxes and target genes in K562 cells. **A**) Efficiency of disruption of selected E-boxes and the spectrum of mutations introduced by individual sgRNAs, demonstrated by TIDE analysis. Size distribution of introduced indels ranged from ≤-30 bp to +2 bp. Colors indicate percentage of sequences with a given indel size. **B**) qRT-PCR analysis of genes adjacent to selected Eboxes upon CRISPR/Cas9 disruption of E-box sequences. For six out of seven E-boxes at least one nearby gene showed significantly decreased expression. Known MYC-regulated genes are underlined; genes essential or at least 4-fold depleted in Brunello screen are in red. Mean and SD of two independent experiments, each performed in triplicate, are shown. **, p<0.01; ***, p<0.001, Student’s t-test. **C**) Cell viability upon disruption of selected E-boxes and knockout of adjacent genes was measured using CellTiter-Glo assay at three timepoints: 0, 48 and 96 hours. Shown are mean values and SD from three independent experiments, each performed in triplicate. *, p<0.05; **, p<<.01; ***, p<0.001; ****, p<0.0001, Student’s t-test. **D**) Luciferase reporter assay for selected E-boxes upon MYC knockdown with shRNA. Decreased luminescence signal was observed for all Eboxes in MYC-shRNA samples vs. NT control. **, p<0.01; ****, p<0.0001, Student’s t-test. Mean and SD of three independent experiments, each performed in triplicate, are shown. **E**) MYC ChIP-qPCR
analysis of MYC binding upon E-box disruption. Cells were infected with sgRNAs targeting selected E-boxes. MYC binding was decreased for chr17_BS377 and for chr10_BS212 but not for chr11_BS79.

Next, we confirmed that for 6 out of 8 selected E-boxes expression of adjacent genes was significantly affected (Figure 4B). In some instances, expression of both genes nearby an E-box was altered, while in others one of the genes was not affected at all or to a lesser extent. Interestingly, for chr17_BS377 we observed strong downregulation of *PFAS* when targeting the essential CACGTG Ebox (sg1), but much weaker effect for the non-essential CGCGTG E-box (sg2 and sg3). For E-box chr2_BS1664 we could not reliably detect expression of the adjacent lncRNAs, neither in control nor CRISPR/Cas9 edited samples. For chr11_BS2113 the closest gene was *PRKRIR* located >50 kb, and we did not observe an impact on its expression. Thus, disruption of E-boxes with CRISPR/Cas9 modulates expression of target genes and can indicate genes regulated by MYC within a given locus.

We further validated the effect on cell growth observed in high-throughput screens for three E-boxes with the strongest effect on expression of adjacent genes: chr10_BS212, chr11_BS79 and chr17_BS377. We conducted growth assays in K562 cells transduced with sgRNAs for E-box disruption and knockout of adjacent genes, and shRNAs for knockdown of the lncRNA PRKCQ-AS1, which was not included in the Brunello screen. For all sgRNAs targeting chosen E-boxes we observed a significant decrease in cell growth, consistent with the results of the screen. We confirmed that within chr17_BS377, only the CACGTG E-box targeted by sg1 is essential for K562 cells growth. Moreover, knockout/knockdown of genes adjacent to each E-box also significantly reduced cell growth, in line with the effect observed in the Brunello screen (Figure 4C). Although both adjacent genes showed decreased expression after disruption of chr10_BS212 and chr11_BS79, only one of each pair was essential for cell growth. This allowed us to pinpoint the MYC targets relevant for the cell growth.

To confirm direct MYC binding and regulation of transcription, we performed a luciferase reporter assay for the three selected E-boxes. We observed decreased luminescent signal for all three MYC binding sites upon MYC knockdown (Figure 4D). This indicates that binding of MYC to these sequences results in MYC-dependent transcription. Moreover, MYC-ChIP in cells transduced with sgRNAs targeting two of the selected E-boxes confirmed decreased MYC binding to chr17_BS377 and chr10_BS212 as compared to non-targeting control sgRNA. Disruption of chr11_BS79 did not affect the strength of MYC binding (Figure 4E).

As an alternative approach, to further validate the importance of MYC binding we utilized dCas9 to block the E-box sequences rather than to disrupt them when using WT Cas9. RT-qPCR in cells transduced with dCas9 and sgRNAs targeting chr10_BS212, chr11_BS79 and chr17_BS377 showed the same pattern of gene expression as for WT Cas9 (Supplementary Figure S3A). In addition, the effect on cell growth for chr11_BS79 and chr17_BS377 was similar with dCas9 and WT Cas9 (Supplementary Figure S3B), while we did not notice change in K562 growth for the sgRNA targeting chr10_BS212. These results further confirm that disturbing MYC binding at these positions is causative for the observed effects on expression of target genes and cell growth.

Altogether, we confirmed that our approach allows for efficient disruption of E-box motifs, which results in decreased MYC binding and affects expression of target genes.

### Implications of the sequence context on E-box functionality

We observed that in some cases with a +1 insertion, the inserted nucleotide did not change the E-box motif (i.e. G added before the last G in the E-box or C inserted after the first C in the E-box). For some sgRNAs the estimated frequency of such DNA edits reached up to ~50% (Supplementary Table S8). Despite apparently not affecting the E-box sequence, we did observe an effect on cell growth and expression of adjacent genes on bulk infected K562 cells. To gain further insights into the E-box grammar, we focused on the E-box CACATG on chromosome 10 (chr10_BS212) with a cut site directly before the last G in the E-box (CACAT*G) and the highest percentage (56%) of +1 insertions with an additional G (CACGTGG).

We successfully established 24 clones of K562 cells with varied mutations and/or WT sequence on all 3 alleles (K562 cells are triploid) (Supplementary Table S9). No significant differences between WT homozygotes and mutants were observed in the expression levels of the nearby PRKCQ gene that was not essential for K562 cells in the Brunello screen. In contrast, expression of the adjacent lncRNA PRKCQ-AS1, whose downregulation negatively affected K562 cell growth, was significantly decreased in +G homozygotes, to a similar extent as in clones with other indels which clearly disrupted the Ebox (Figure 5A). This indicated that even though the E-box motif sequence *per se* was not changed, its functionality was affected.

**Figure 5.**
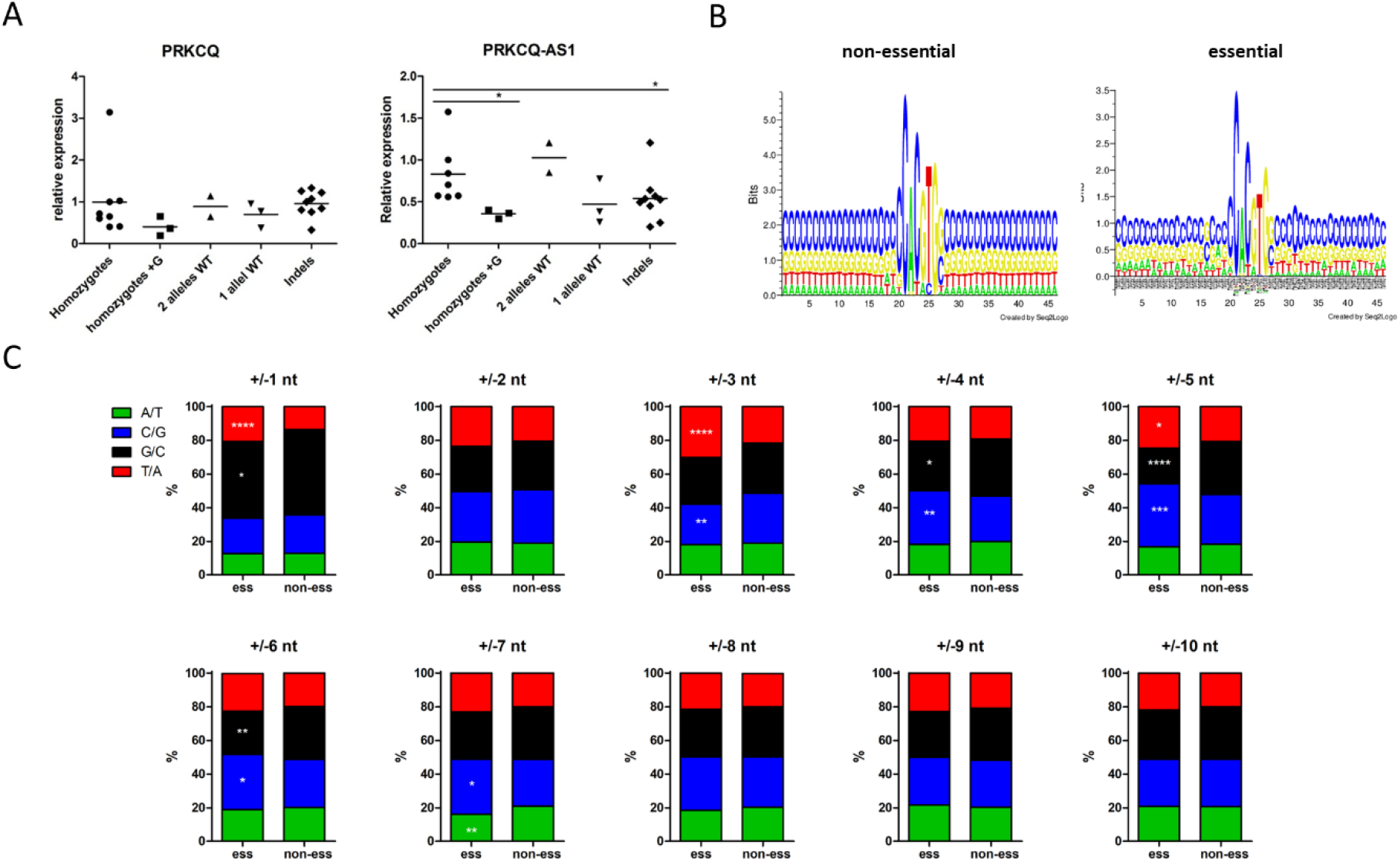
Grammar of the E-box sequence context. **A**) Expression of two genes adjacent to an E-box on chromosome 10 (chr10_BS212) was examined in monoclonal cell lines derived from K562 cells transduced with an sgRNA targeting this E-box. A spectrum of clones with various modifications of the E-box were obtained, including homozygotes with the +G insertion after E-box which seemingly did not change the E-box sequence. *, p<0.05, Kruskal-Wallis test with Dunn’s post test. **B**) Sequence logo (created using Seq2Logo) of the E-boxes and 20 nt flanking sequences for non-essential (left) and essential (right) E-boxes. **C**) Frequency of up to 10 nt upstream/downstream flanking essential and non-essential E-boxes. Since E-boxes are (quasi)palindromic and can be read on either strand, G at +1 equals C at −1 etc. *, p<0.05; **, p<0.01; chi-square goodness of fit test.

This prompted us to look at the sequences flanking the E-boxes. Analysis of the nucleotide frequency 20 nt upstream and downstream of non-essential E-boxes showed uniform distribution of nucleotides. In contrast, there were marked differences at particular positions flanking essential E-boxes (Figure 5B). Statistical analysis of 10 nt upstream/downstream revealed that certain nucleotides were significantly over- or underrepresented in the neighborhood of essential E-boxes (Figure 5C). Since E-boxes are (quasi)palindromic and can be read on either strand, G at +1 equals C at −1 etc. The strongest bias in nucleotide composition was observed at positions +/-1 nt, and 3-7 nt. In particular, we observed that G immediately after or C immediately before essential E-boxes was unfavored. Thus, we speculate that the immediate context of E-boxes might affect their functionality. This could explain why such changes without apparent disruption of the E-box affected expression of adjacent genes and cell viability.

## Discussion

Despite decades of research, MYC still evades full comprehension and therapeutic targeting. To understand the mechanisms underlying cancer cell addiction to MYC, it is essential to determine the crucial genes regulated by this TF. Therefore, the aim of this study was to identify on a genome-wide scale functional MYC binding sites and corresponding target genes essential for cancer cells growth. To this end, we have established a novel CRISPR/Cas9-based tool to disrupt MYC-bound E-boxes.

Extensive validation of our approach confirmed efficient E-box disruption, decreased expression of adjacent genes and reduced MYC binding upon CRISPR/Cas9 editing of selected Eboxes. No change in MYC binding for the E-box on chr11 can be potentially explained by the presence of another, non-essential E-box, chr11_BS79_CACGCG ~100 bp downstream of the analyzed E-box chr11_BS79_CGCGTG (Supplementary Figure S5B). Resolution of ChIP-PCR with DNA fragments of ca. 500 bp does not allow to distinguish such close binding sites. Finally, using individual sgRNAs we confirmed that E-box disruption or knockout of adjacent genes significantly decreased K562 cell proliferation, in line with the results of the screens. However, we observed some discrepancies using a parallel dCas9 approach. We validated the effect on cell growth for two out of three E-boxes but for chr10_BS212 we did not observe decreased proliferation with dCas9, despite a similar effect on expression of two adjacent genes *PRKCQ* and *PRKCQ-AS1*. Similar inconsistencies have also been reported for targeting p53 binding sites with WT Cas9 and dCas9 with an overall limited overlap^31^ and this phenomenon requires further investigation.

Notably, using our strategy it was possible to determine which E-boxes are essential for cell viability and identify relevant regulated target genes. This is important since we demonstrated that not all E-boxes in a binding site affected cell growth and target gene expression. Similarly, not all genes adjacent to an E-box responded to E-box disruption and were crucial for cancer cells. In our screen, 68-80% of essential E-boxes were not localized near essential genes, but this might be an underestimation as we checked only the first TSS within 50 kb up- and downstream of each E-box. Moreover, due to chromatin organization, the relevant target may be much further away. In addition, MYC also regulates non-coding RNAs, which were not included in this general analysis, yet might be essential. Combination of the MYC-CRISPR screen with single cell RNA sequencing would add another layer of information to this experimental setup and allow direct identification of genes responding to E-box disruption.

Essential genes identified in the Brunello screen showed a substantial overlap, and 77% of them belong to the panel of pan-essential genes^32^. GO and GSEA analyses revealed processes common for all cell lines which indicates that different types of cancers rely on the same factors for their growth^33^. On the other hand, the overlap between essential E-boxes was very limited and analysis of adjacent genes revealed a bigger spectrum of cell type-specific processes. This observation is in line with the fact that MYC acts within the pre-defined transcriptional landscape which varies between cell types and developmental stages^20,34,35^.

A recent study identified 1344 MYC-dependent genes (log2FC <-0.58, p<0.05) in K562 cells using SLAM-seq upon MYC disruption^36^. 1035 of these genes had an E-box within 50 kb that was included in the MYC-CRISPR library. Of these, 284 genes were localized near an E-box that was at least 1.5-fold depleted in our screen in K562 cells (Supplementary Figure S6A, 6B). This limited overlap may be due to the fact that in our study we focused on MYC targets which were essential for K562 cells growth, while SLAM-seq included all targets. On the other hand, due to sgRNA design in our screen we might have missed some E-boxes relevant for target genes. This highlights the need for integration of multiple approaches for identification of essential MYC targets.

Interestingly, our study provided also some novel insights into the grammar of E-boxes and their surrounding sequences. We observed significantly different frequencies of specific nucleotides at certain positions in essential vs non-essential E-boxes. This observation provides novel indications about E-box functionality and demonstrates the usefulness of high-throughput CRISPR/Cas9 mutagenesis for studying TF binding sites. Previous studies showed that DNA flexibility and structure determined by the flanking sequences impacts binding of TFs^37^, including closely related TFs from the bHLH family: MYC, MAX and MAD^38,39^. Phylogenetic comparisons revealed strong sequence conservation of E-boxes and also their flanking regions among species^40^. Moreover, it was recently reported that MYC is first engaged in open chromatin regions *via* non-specific binding, while recognition of specific sequences stabilizes binding of MYC to DNA and promotes its transcriptional activity^41^.

A limitation of our study is that we were not able to target all E-boxes within the MYC-bound loci. This was due to either lack of PAM sequence nearby (35% of E-boxes not included in our library) or strong off-target activity of designed sgRNAs (only 4% of E-boxes not targeted). This could be overcome with a complementary approach using variant Cas9 nucleases with different PAM requirements.

A potential flaw in our approach could be the fact that other bHLH proteins can bind to E-boxes and affect transcription^42^. Therefore, effect of E-box disruption might be also related to other interactors. However, several findings strongly suggest that MYC is involved: 1) we focused on validated MYC binding sites (based on available MYC ChIP-seq data); 2) luciferase reporter assay for selected Eboxes showed decreased transcription after MYC knockdown; 3) ChIP confirmed that E-box disruption reduced MYC binding. Altogether, this indicates that although we cannot exclude that other factors may also be involved, MYC binding is crucial for the activity of the studied E-boxes.

In summary, the combined high-throughput screens using the MYC-CRISPR library targeting E-boxes and the Brunello library for gene knockout is a first powerful tool for genome-wide identification of MYC-dependent vulnerabilities in cancer cells. This well-validated novel approach allows for the identification of essential MYC binding sites and the regulated target genes which can serve as a basis for novel therapeutic approaches in MYC-dependent cancers. The broad design enables studies in a variety of normal and cancer cell types and determination of common as well as cell-type specific targets.

## MATERIALS AND METHODS

### Cell lines

K562, ST486 and HepG2, MCF7, HEK293T cells were cultured in RPMI or DMEM (Lonza, Basel, Switzerland) respectively, supplemented with 10% fetal bovine serum (Sigma-Aldrich, Saint Louis, MO, US), 2mM L-glutamine and 1% penicillin/streptomycin (Biowest, Nuaillé, France). In addition, medium for MCF7 contained 1x NEAA (Gibco, Waltham, MA US).

### Plasmids

The lentiCRISPR v2 (#52961)^43^ and lentiCRISPR v2-dCas9 (#112233)^44^ vectors were purchased from Addgene (Watertown, MA, US).

### Design and cloning of the MYC-CRISPR library

We utilized publicly available MYC**-**ChIP-seq data from MCF7, K562 and HepG2 cells^27^ and BL cell lines. E-box motifs were identified (canonical CACGTG, non-canonical CACATG/CATGTG and CACGCG/CGCGTG) and all sgRNAs targeting these E-boxes were designed based on the presence of the PAM sequence (NGG or CCN) using an in-house Python script (https://github.com/tomaszwozniakihg/cas9_search_tool). The resulting sgRNAs were checked for off-target binding using the CAS-OT script^45^. The library also included 1 000 non-targeting sgRNAs as a negative control, and four sgRNAs against MYC as a positive control, all from the Brunello library. A list of all sgRNA oligonucleotides is provided in Supplementary Table S1. Oligonucleotides containing the 20 nt sgRNA sequences flanked by the sequence from the lentiCRISPR_v2 vector were synthesized by Twist Bioscience (San Francisco, CA, US). Oligo amplification and library cloning was performed as described previously^46^. The MYC-CRISPR library was deposited in Addgene (#173195). Brunello library targeting all human protein-coding genes^28^ was purchased from Addgene (#73179). Quality of the MYC-CRISPR and Brunello libraries was verified by next-generation sequencing on Illumina platform (BGI, Hong-Kong). Individual sgRNAs (Supplementary Table S10) were cloned into the lentiCRISPRv2_puro and lentiCRISPRv2-dCas9 vectors with T4 DNA ligase (Invitrogen, Carlsbad, CA, US).

### CRISPR/Cas9 screens

Lentiviral particles were produced in HEK293T cells using 2^nd^ generation packaging plasmids. 78 million cells for the MYC-CRISPR library and 130 million cells for the Brunello library were transduced in duplicate with the amount of virus that results in ~30% transduced cells. 4 mg/ml polybrene was added and cells were spun down in plates (33°C, 1000x g, 2h). After four days of selection with puromycin (T0) part of the cells was collected for DNA isolation. Remaining cells were further cultured for 20 population doublings. At each passage, the amount of cells corresponding to a 500x coverage (24 million for MYC-CRISPR, 38 million for Brunello library) were cultured in RPMI medium with 1 μg/ml puromycin and collected at the final timepoint (T1).

### NGS and data analysis

sgRNA sequences from the MYC-CRISPR and Brunello library plasmids and from DNA from collected cells were amplified in PCR reaction as described previously^46^ (Supplementary Table S11). NGS was performed on Illumina X-Ten platform at BGI (Hong-Kong). Number of reads obtained for each sample is given in Supplementary Table S12. For sgRNA enumeration, raw reads were processed with a Python script^46^. sgRNA counts were then used for DeSeq2 analysis with the CRISPRAnalyzeR tool (http://www.crispr-analyzer.org)^47^. Adjusted p-value 0.001 was used as a cut-off for identification of significantly depleted or enriched E-boxes (MYC-CRISPR library) and genes (Brunello library).

### Validation of the approach

To confirm CRISPR/Cas9 disruption of selected E-boxes, genomic regions of ~500-800 bp flanking E-box sequences were amplified by PCR and analyzed with TIDE calculator (https://tide-calculator.nki.nl)^48^. Effect of E-box disruption on expression of adjacent genes was determined by RT-qPCR. Growth assay was performed using CellTiter-Glo^®^ Luminescent Cell Viability Assay (Promega, Madison, WI, USA). To confirm that sequences containing selected E-boxes drive transcription in a MYC-dependent way, we conducted luciferase reporter assay upon MYC knockdown. See Supplementary Information for detailed description.

### Chromatin immunoprecipitation

10 M of K562 cells infected with sgRNAs targeting selected E-box sequences were fixed to crosslink DNA with chromatin-associated proteins according to the Active Motif protocol. Chromatin immunoprecipitation with anti-MYC antibody (sc-764, Santa Cruz, Dallas, TX, US) followed by qPCR was performed by Active Motif (La Hulpe, Belgium).

### Gene Ontology and Gene Set Enrichment Analysis

For each E-box, the nearest genes with transcription start site (TSS) within 50kb both upstream and downstream were retrieved using Galaxy. Gene ontology analysis was conducted using DAVID Functional Annotation Tool v6.8b^49,50^ Pre-ranked gene set enrichment analysis (GSEA)^51,52^ was performed on log2 fold change values using Hallmark (H) and curated (C2) gene sets v7.4.

### Statistical analysis

To determine the statistical significance of results from CellTiter-Glo, RT-qPCR and luciferase assay Student’s t-test was applied. Enrichment of essential genes near essential E-boxes was assessed by the chi-square goodness of fit test. E-box flanking sequences were analyzed with the chi-square goodness of fit test and expression of genes for different K562 clones was analyzed using Kruskal-Wallis test with Dunn’s post test. All statistical analyses were performed in GraphPad Prism, p value <0.05 was considered significant.

## AUTHORS’ CONTRIBUTIONS

AD-K planned and supervised the project and acquired funding; MK, MŻ, MP, MEK, WS, WŁ, IZ-S and AD-K performed the experiments; TW designed the MYC-CRISPR library sgRNAs; data were analyzed by MK, MP, WŁ, NR and AD-K; NR, JK and AvdB contributed to project conceptualization; MK and AD-K prepared the figures and wrote the manuscript. All authors read and approved the manuscript.

## ACKNOWLEDGEMENTS

We would like to thank Dr. Jeroen Guikema from University Medical Center Amsterdam, the Netherlands, and Dr. Maciej Giefing from the Institute of Human Genetics, Poland, for inspiring discussions.

Figures: 2A and 2B were created using BioRender.

## DATA AVAILABILITY

Script used to generate MYC-CRISPR library is deposited on github: https://github.com/tomaszwozniakihg/cas9_search_tool

MYC-CRISPR library was deposited in Addgene (#173195).

