## Supplementary Figures, Tables and Methods for "CRISPR/Cas9 screen for genome-wide interrogation of essential MYC binding sites in cancer cells"

### SUPPLEMENTARY INFORMATION

#### MATERIALS AND METHODS

##### Cell lines

The chronic myeloid leukemia cell line K562 (ordered from DSMZ, Braunschweig, Germany) and Burkitt lymphoma cell line ST486 (ATCC, Manassas, VA, US) were maintained in Roswell Park Memorial Institute 1640 medium (Lonza, Basel, Switzerland) supplemented with 10-20% fetal bovine serum (Sigma-Aldrich, Saint Louis, MO, US) 2mM L-glutamine (Biowest, Nuaille, France), and 1% penicillin/streptomycin (Biowest) in a 5% $CO_2$ incubator at 37°C. Hepatocellular carcinoma cell line HepG2 (DSMZ), breast cancer cell line MCF7 (ECACC, Porton Down, United Kingdom) and HEK293T cell line (DSMZ) used for lentiviral particles production were cultured in low glucose Dulbecco's Modified Eagle's Medium (Lonza) supplemented as described above. In addition, medium for MCF7 cells was supplemented with 1x NEAA (Gibco, Waltham, MA US). Mycoplasma tests were routinely performed and confirmed that the cells were not contaminated.

**Plasmids**

The lentiCRISPR v2 (#52961)[1] and lentiCRISPR v2-dCas9 (#112233)[2] vectors were purchased from Addgene (Watertown, MA, US). The plasmids contains the coding sequence of *S. pyogenes* (d)Cas9 and a cloning site for the sgRNA, as well as a puromycin resistance gene allowing selection of transduced cells. pGreenpuro lentivectors (SBI, Mountain View, CA, USA) with MYC shRNAs and scrambled control, a kind gift of prof. Anke van den Berg[3], were used for experiments with MYC knockdown.

**Design of the MYC-CRISPR library**

To design a library of single guide RNA (sgRNA) for comprehensive genome-wide targeting of MYC binding sites in cancer cells, we utilized publicly available MYC-ChIP-seq data from MCF7, K562 and HepG2 cells[4] and Burkitt lymphoma cell lines[5]. Genomic coordinates (hg19) of MYC-ChIP peaks were obtained using UCSC Table Browser and from the published data[5]. Genomic intervals were concatenated, merged, and DNA sequences were retrieved using Galaxy[6]. To avoid targeting coding exons, which could have resulted in disruption of the protein apart from the E-box, sequences overlapping with coding exons (based on Ensembl hg19 annotation) were removed. In the resulting sequences, E-box motifs were identified (canonical CACGTG, non-canonical CACATG/CATGTG and CACGCG/CGCGTG) and all sgRNAs targeting these E-boxes were designed based on the presence of the PAM sequence (NGG or CCN) using an in-house Python script (https://github.com/tomaszwozniakihg/cas9_search_tool). The resulting sgRNAs were checked for off-target binding using the CAS-OT script[7]. Only sgRNAs with at least three mismatches to the potential off-targets or at least two mismatches including at least one in the seed region (nt 9-20) were retained. The library also included 1,000 non-targeting sgRNAs as a negative control, and four sgRNAs against MYC as a positive control, all from the Brunello library. A list of all sgRNA oligonucleotides is provided in Table S1.

**Cloning of the MYC-CRISPR library and amplification of the Brunello library**

Oligonucleotides containing the 20 nt sgRNA sequences flanked by the sequence from the lentiCRISPR_v2 vector TTTCTTGGCTTTATATATCTTGTGGAAAGGACGAAACACCG [20 nt sgRNA]

GTTTTAGAGCTAGAAATAGCAAGTTAAAATAAGGCTAGTCCGT were synthesized by Twist Bioscience (San Francisco, CA, US). 2 ng of oligonucleotide pool was amplified with the oligo-F and oligo-R primers (Supplementary Table S11) using the NEBNext High-Fidelity Master Mix (New England Biolabs, Ipswich, MA, US) in 20 x 25 µl PCR reactions. PCR program: 98°C 30 sec; (98°C 10 sec; 63°C 10 sec; 72°C 15 sec) x 7 cycles; 72°C 2 min. PCR product was purified using QIAquick PCR Purification Kit (Qiagen, Hilden, Germany) and then extracted from an agarose gel with QIAquick Gel Extraction Kit (Qiagen). LentiCRISPRv2_puro vector was digested with the BsmBI restriction enzyme (New England Biolabs) and purified from an agarose gel. sgRNA MYC-CRISPR library was cloned into lentiCRISPRv2_puro vector using the circular polymerase extension cloning (CPEC) method as described previously[8]. Briefly, 20 CPEC reactions were performed, each using 100 ng digested vector, 10.8 ng amplified oligos and NEBNext High-Fidelity Master Mix. PCR program: 98°C 30 sec; (98°C 10 sec; 72°C 7 min) x 5 cycles; 72°C 5 min. All PCR reactions were pooled and purified by isopropanol precipitation. 300 ng of the CPEC product was used for transformation of electrocompetent Endura cells (Lucigen, Middleton, WI, USA) according to the manufacturer's protocol. Fourteen electroporations were performed giving in total ~7.9 million colonies and resulting in ~170x coverage of the library. Bacteria were spread on 245x245 agar plates and grown for 14 h at 37°C. Colonies were scraped off the plates and plasmid DNA was isolated using Plasmid Plus Maxi Kit (Qiagen). The MYC-CRISPR library was deposited in Addgene (#173195). Brunello library targeting all human protein-coding genes[9] was purchased from Addgene (#73179). 50 ng of the library was electroporated into Endura cells. Four electroporations were performed resulting in >2,500x coverage. Quality of the MYC-CRISPR and Brunello libraries was verified by next-generation sequencing on Illumina platform (BGI, Hong-Kong).

**Cloning of individual sgRNA constructs**

For cloning of individual sgRNAs (Supplementary Table S10) into the lentiCRISPRv2_puro and lentiCRISPRv2-dCas9 vectors, sense and antisense oligos containing overhangs compatible with BsmBI sticky ends were synthesized by Genomed (Warsaw, Poland). Oligonucleotides were diluted in annealing buffer (10mM Tris-HCl pH8, 1mM EDTA pH8, 50mM NaCl). Annealing was performed in a thermocycler under conditions: 95°C 5min, 95°C (-1°C/cycle) x 70 cycles. Annealed sgRNAs were ligated into the lentiCRISPRv2 and lentiCRISPRv2-dCas9 vectors digested with BsmBI restriction enzyme at 1:5 vector:insert molar ratio with T4 DNA ligase (Invitrogen, Carlsbad, CA, US). 1 µl of ligation reaction was transformed into JM109 competent cells (Promega, Madison, WI, USA). Plasmid DNA from single colonies was isolated using Plasmid Plus Maxi Kit (Qiagen). Sequences of individual sgRNA constructs were confirmed by Sanger sequencing (Genomed).

**Generation of lentiviral particles**

For large scale production of lentiviral particles ~7.5 million HEK293T cells were plated in a T75 flask one day prior to transfection. Next day, ~80% confluent cells were transfected with packaging plasmids psPAX (11.2 µg) and pMD2.G (7.5 µg), and MYC-CRISPR or Brunello library plasmid (15 µg) using lipofectamine 2000 reagent (Invitrogen). One day after transfection medium was replaced with 7.5 ml DMEM supplemented with 10% FBS. Two and three days post-transfection Brunello and MYC-CRISPR lentiviral supernatants were filtered through 0.45 µm filter and stored at -80°C. For small scale production of lentiviral particles 1 million HEK293T cells were plated per well on a 6-well plate and transfected the next day with packaging plasmids psPAX (1.5 µg) and pMD2.G (1 µg), and lentiCRISPRv2 plasmid (2 µg) using calcium phosphate transfection method (Invitrogen). One day after transfection medium was replaced with 1.1 ml DMEM supplemented with 10% FBS and lentiviral supernatant was collected as described above.

**Determination of virus titer and cell transduction for screening**

1.8-2.7 million cells were plated per well in a 12-well plate and transduced with different amounts of virus. 4 mg/ml polybrene was added and cells were spun down in plates (33°C, 1000x g, 2h). After spinfection, additional 1 ml of medium was added. 24 h after transduction cells were washed and plated out in four wells for each condition, two wells with 0.3-3 µg/ml puromycin and two wells without puromycin. Medium with puromycin was changed after three days. After four days of selection, cells were counted and the percentage of surviving cells relative to cells not treated with puromycin was calculated. From this we determined the amount of virus resulting in ~30% surviving cells, which implies that approximately 85% of transduced cells were infected by a single virus.

~78 million cells for the MYC-CRISPR library and ~130 million cells for the Brunello library were transduced in duplicate with the amount of virus that results in ~30% transduced cells, in the same conditions as described above. After four days of selection with puromycin (T0) part of the cells was collected for DNA isolation. Remaining cells were further cultured for 20 population doublings. At each passage, the amount of cells corresponding to a 500x coverage (24 million for MYC-CRISPR, 38 million for Brunello library) were cultured in RPMI medium with 0.1-1 µg/ml puromycin and collected at the final timepoint (T1).

**Preparation of libraries for next-generation sequencing of the plasmid pool or genomic DNA**

sgRNA sequences from the MYC-CRISPR and Brunello library plasmids were amplified in PCR reaction using High Fidelity MasterMix 2x (New England Biolabs) and primers containing Illumina adaptors, an 8 nt barcode specific for each library (reverse primer) and a variable length (9-18 nt) stagger sequence to increase library complexity (forward primer)[10] (Supplementary Table S11). PCR products were pooled and purified using QIAquick PCR Purification Kit (Qiagen). Amplicons were analyzed on agarose gel and then extracted using QIAquick Gel Extraction Kit (Qiagen). Quality check and quantification of libraries were performed by qPCR using KAPA Library Quantification Kit (Roche, Basel, Switzerland).

Genomic DNA collected from cells infected with MYC-CRISPR and Brunello libraries was isolated using GENTRA Puregene Kit (Qiagen). PCR was performed to amplify the sgRNA sequences integrated in genomic DNA. DNA from 23 million cells (MYC-CRISPR) and 38 million cells (Brunello) was amplified in 80-120 (MYC-CRISPR) and 130-210 (Brunello) individual 50 µl PCR reactions per sample with 3-3.5 µg DNA input as described above. PCR products were purified, analyzed on agarose gel, pooled based on band intensities, and extracted from gel as described above. Determination of quality and quantification of libraries was performed by qPCR (Kapa Library Quantification Kit).

**NGS and data analysis**

NGS was performed on Illumina X-Ten platform at BGI (Hong-Kong). Reads were trimmed to remove adaptor sequences and split based on barcodes for individual samples. Number of reads obtained for each sample is given in Supplementary Table S12. For sgRNA enumeration, raw reads were processed with a Python script[10]. Only sgRNAs with no mismatches were counted. sgRNA counts were then used for DeSeq2 analysis with the CRISPRAnalyzeR tool (http://www.crispr-analyzer.org)[11]. Adjusted p-value 0.001 was used as a cut-off for identification of significantly depleted or enriched E-boxes (MYC-CRISPR library) and genes (Brunello library).

**Determination of CRISPR/Cas9 editing efficiency by TIDE**

To confirm CRISPR/Cas9 disruption of selected E-boxes, DNA was isolated from K562 cells at day 7 after transduction with individual sgRNAs targeting the selected E-boxes. Genomic regions of ~500-800 bp flanking E-box sequences were amplified by PCR (primer sequences in Supplementary Table S11). Amplicons were sequenced using the Sanger method (Genomed) and analyzed with TIDE calculator (https://tide-calculator.nki.nl)[12], using indel size range of 50.

**RNA isolation and qRT-PCR**

Total RNA was isolated from K562 cells using Quick-RNA™ Miniprep Kit (Zymo Research, Irvine, CA, US). cDNA was synthesized from 500 ng RNA using QuantiTect® Reverse Transcription Kit (Qiagen). qPCR on 5 ng cDNA was performed on CFX96 Touch qPCR System (BioRad, Hercules, CA, US) using PowerUp SYBR Green Master Mix (Applied Biosystems, Waltham, MA US). Expression was normalized to TBP. All experiments were conducted in two independent biological replicates, each with three technical replicates.

**Growth assay**

Growth assay was performed using CellTiter-Glo® Luminescent Cell Viability Assay (Promega). $1*10^3$ K562 cells infected with sgRNA lentiviral constructs were plated in triplicate on 96-well plates. 100 ul Cell-Titer Glo reagent diluted 1:2 in PBS was added per well after 1h (baseline level), 48h and 96h. The luminescent signal was measured using a GloMax microplate reader (Promega). Experiments were performed in three independent biological replicates. Growth rate was calculated at 48h and 96h relative to the 1h measurement.

**Chromatin immunoprecipitation assay**

10 M of K562 cells infected with sgRNAs targeting selected E-box sequences were fixed to crosslink DNA with chromatin-associated proteins according to the Active Motif protocol. Briefly, cells were fixed for 20 min by adding 1/10 volume of 37% formaldehyde solution and then neutralized by adding 1/20 volume of 2.5 M glycine. Next, K562 cells were washed using PBS-Igepal, snap-frozen on dry ice and stored at -80$^{o}$C. Chromatin immunoprecipitation with anti-MYC antibody (sc-764, Santa Cruz, Dallas, TX, US) followed by qPCR was performed by Active Motif (La Hulpe, Belgium).

**Luciferase reporter assay**

To confirm that sequences containing selected E-boxes drive transcription in a MYC-dependent way, we conducted luciferase reporter assay. MYC binding sites as defined by peaks from MYC-ChIP-seq data were amplified by PCR from DNA of K562 cells. For chr11_BS79 the amplified sequence was shorter to omit two other non-essential E-boxes present in the peak. Primers contained sequences to create overhangs for XhoI and SacI restriction sites (Supplementary Table S11) for cloning into the pGL4.23 vector, upstream of the firefly luciferase gene under minimal promoter (#E8411, Promega). K562 cells were transduced with MYC shRNAs or scrambled control. After selection with puromycin for 4 days, 1x$10^5$ cells were co-transfected using Lipofectamine LTX with 500 ng pGL4.23 vector with E-box constructs and 5 ng pRL-SV40 vector containing Renilla luciferase (#E2231, Promega). 24h after transfection cells were lysed and luminescence was measured using Dual Luciferase Reporter Assay on GloMax (Promega). Firefly luminescence was normalized to Renilla. The experiments were performed in three independent biological replicates, each with triplicate transfection.

**Gene Ontology and Gene Set Enrichment Analysis**

For each E-box, the nearest genes with transcription start site (TSS) within 50 kb both upstream and downstream were retrieved using Galaxy. Gene ontology analysis was conducted using DAVID Functional Annotation Tool v6.8b[13,14] on essential genes from Brunello screen and at least two-fold depleted genes within 50 kb of essential E-boxes from MYC-CRISPR screen. Pre-ranked gene set enrichment analysis (GSEA)[15,16] was performed on log2 fold change values of all genes in the Brunello library and of genes near at least two-fold depleted E-boxes. Hallmark (H) and curated (C2) gene sets v7.4 were used for analysis.

### SUPPLEMENTARY TABLES

**Table S6.** Top 50 enriched gene sets among genes from Brunello library.

| | K562 | | | ST486 | | | HepG2 | | | MCF7 | | |
|---|---|---|---|---|---|---|---|---|---|---|---|---|
| **GENE SETS** | **RANK** | **NES** | **FDR q-val** | **RANK** | **NES** | **FDR q-val** | **RANK** | **NES** | **FDR q-val** | **RANK** | **NES** | **FDR q-val** |
| KEGG_RIBOSOME | 1 | -2.28 | 0.0000 | 3 | -1.81 | 0.0000 | 1 | -2.02 | 0.0000 | 8 | -3.14 | 0.0000 |
| REACTOME_EUKARYOTIC_TRANSLATION_ELONGATION | 3 | -2.24 | 0.0000 | 2 | -1.82 | 0.0000 | 6 | -1.98 | 0.0000 | 5 | -3.16 | 0.0000 |
| REACTOME_RRNA_PROCESSING | 6 | -2.19 | 0.0000 | 7 | -1.80 | 0.0000 | 5 | -1.98 | 0.0000 | 2 | -3.29 | 0.0000 |
| REACTOME_EUKARYOTIC_TRANSLATION_INITIATION | 10 | -2.17 | 0.0000 | 1 | -1.82 | 0.0000 | 8 | -1.97 | 0.0000 | 3 | -3.22 | 0.0000 |
| REACTOME_TRANSLATION | 5 | -2.20 | 0.0000 | 9 | -1.80 | 0.0000 | 10 | -1.94 | 0.0000 | 1 | -3.33 | 0.0000 |
| REACTOME_RESPONSE_OF_EIF2AK4_GCN2_TO_AMINO_ACID_DEFICIENCY | 8 | -2.19 | 0.0000 | 8 | -1.80 | 0.0000 | 2 | -1.99 | 0.0000 | 7 | -3.14 | 0.0000 |
| WP_CYTOPLASMIC_RIBOSOMAL_PROTEINS | 2 | -2.24 | 0.0000 | 11 | -1.79 | 0.0000 | 4 | -1.98 | 0.0000 | 9 | -3.11 | 0.0000 |
| REACTOME_NONSENSE_MEDIATED_DECAY_NMD | 4 | -2.22 | 0.0000 | 4 | -1.80 | 0.0000 | 11 | -1.94 | 0.0000 | 10 | -3.10 | 0.0000 |
| KEGG_SPLICEOSOME | 11 | -2.16 | 0.0000 | 5 | -1.80 | 0.0000 | 7 | -1.97 | 0.0000 | 17 | -2.96 | 0.0000 |
| REACTOME_MRNA_SPLICING | 15 | -2.13 | 0.0000 | 6 | -1.80 | 0.0000 | 9 | -1.95 | 0.0000 | 14 | -3.06 | 0.0000 |
| REACTOME_INFLUENZA_INFECTION | 13 | -2.13 | 0.0000 | 10 | -1.79 | 0.0000 | 18 | -1.92 | 0.0000 | 4 | -3.17 | 0.0000 |
| REACTOME_SELENOAMINO_ACID_METABOLISM | 9 | -2.19 | 0.0000 | 12 | -1.78 | 0.0000 | 12 | -1.94 | 0.0000 | 13 | -3.07 | 0.0000 |
| REACTOME_SRP_DEPENDENT_COTRANSLATIONAL_PROTEIN_TARGETING_TO_MEMBRANE | 7 | -2.19 | 0.0000 | 37 | -1.74 | 0.0000 | 3 | -1.99 | 0.0000 | 12 | -3.07 | 0.0000 |
| REACTOME_PROCESSING_OF_CAPPED_INTRON_CONTAINING_PRE_MRNA | 17 | -2.12 | 0.0000 | 21 | -1.76 | 0.0000 | 15 | -1.93 | 0.0000 | 6 | -3.15 | 0.0000 |
| REACTOME_REGULATION_OF_EXPRESSION_OF_SLITS_AND_ROBOS | 12 | -2.14 | 0.0000 | 23 | -1.76 | 0.0000 | 21 | -1.91 | 0.0000 | 11 | -3.08 | 0.0000 |
| REACTOME_ACTIVATION_OF_THE_MRNA_UPON_BINDING_OF_THE_CAP_BINDING_COMPLEX_AND_EIFS_AND_SUBSEQUENT_BINDING_TO_43S | 20 | -2.11 | 0.0000 | 14 | -1.78 | 0.0000 | 13 | -1.94 | 0.0000 | 21 | -2.92 | 0.0000 |
| WP_MRNA_PROCESSING | 22 | -2.10 | 0.0000 | 15 | -1.77 | 0.0000 | 17 | -1.92 | 0.0000 | 25 | -2.86 | 0.0000 |
| HALLMARK_MYC_TARGETS_V1 | 18 | -2.12 | 0.0000 | 26 | -1.76 | 0.0000 | 22 | -1.91 | 0.0000 | 15 | -2.98 | 0.0000 |
| BILANGES_SERUM_AND_RAPAMYCIN_SENSITIVE_GENES | 16 | -2.12 | 0.0000 | 24 | -1.76 | 0.0000 | 19 | -1.92 | 0.0000 | 23 | -2.86 | 0.0000 |
| REACTOME_RRNA_MODIFICATION_IN_THE_NUCLEUS_AND_CYTOSOL | 19 | -2.11 | 0.0000 | 17 | -1.76 | 0.0000 | 23 | -1.91 | 0.0000 | 30 | -2.83 | 0.0000 |
| REACTOME_MRNA_SPLICING_MINOR_PATHWAY | 26 | -2.07 | 0.0000 | 13 | -1.78 | 0.0000 | 20 | -1.91 | 0.0000 | 31 | -2.82 | 0.0000 |
| REACTOME_CELLULAR_RESPONSE_TO_STARVATION | 27 | -2.07 | 0.0000 | 16 | -1.77 | 0.0000 | 36 | -1.89 | 0.0000 | 18 | -2.96 | 0.0000 |
| REACTOME_MITOCHONDRIAL_TRANSLATION | 21 | -2.11 | 0.0000 | 20 | -1.76 | 0.0000 | 35 | -1.89 | 0.0000 | 22 | -2.87 | 0.0000 |
| CHNG_MULTIPLE_MYELOMA_HYPERPLOID_UP | 14 | -2.13 | 0.0000 | 27 | -1.76 | 0.0000 | 29 | -1.90 | 0.0000 | 51 | -2.68 | 0.0000 |
| REACTOME_CHROMOSOME_MAINTENANCE | 28 | -2.06 | 0.0000 | 42 | -1.73 | 0.0000 | 31 | -1.90 | 0.0000 | 35 | -2.79 | 0.0000 |
| REACTOME_DNA_REPLICATION | 39 | -2.04 | 0.0000 | 41 | -1.73 | 0.0000 | 39 | -1.88 | 0.0000 | 20 | -2.93 | 0.0000 |
| REACTOME_TELOMERE_MAINTENANCE | 29 | -2.06 | 0.0000 | 47 | -1.73 | 0.0000 | 27 | -1.90 | 0.0000 | 42 | -2.74 | 0.0000 |
| REACTOME_SIGNALING_BY_ROBO_RECEPTORS | 25 | -2.07 | 0.0000 | 63 | -1.71 | 0.0000 | 40 | -1.88 | 0.0000 | 19 | -2.95 | 0.0000 |
| REACTOME_TRANSCRIPTION_COUPLED_NUCLEOTIDE_EXCISION_REPAIR_TC_NER | 24 | -2.08 | 0.0000 | 35 | -1.74 | 0.0000 | 16 | -1.92 | 0.0000 | 84 | -2.57 | 0.0000 |
| REACTOME_SNRNP_ASSEMBLY | 46 | -2.02 | 0.0000 | 30 | -1.75 | 0.0000 | 48 | -1.85 | 0.0000 | 46 | -2.73 | 0.0000 |
| TIEN_INTESTINE_PROBIOTICS_6HR_UP | 23 | -2.08 | 0.0000 | 62 | -1.71 | 0.0000 | 32 | -1.90 | 0.0000 | 63 | -2.64 | 0.0000 |
| REACTOME_DUAL_INCISION_IN_TC_NER | 41 | -2.03 | 0.0000 | 39 | -1.74 | 0.0000 | 14 | -1.94 | 0.0000 | 108 | -2.51 | 0.0000 |
| REACTOME_DNA_REPLICATION_PRE_INITIATION | 66 | -1.99 | 0.0000 | 72 | -1.70 | 0.0001 | 41 | -1.87 | 0.0000 | 36 | -2.78 | 0.0000 |
| REACTOME_DNA_STRAND_ELONGATION | 38 | -2.04 | 0.0000 | 31 | -1.75 | 0.0000 | 24 | -1.91 | 0.0000 | 124 | -2.48 | 0.0000 |
| WP_DNA_REPLICATION | 42 | -2.03 | 0.0000 | 60 | -1.71 | 0.0000 | 28 | -1.90 | 0.0000 | 89 | -2.56 | 0.0000 |
| REACTOME_HIV_TRANSCRIPTION_ELONGATION | 34 | -2.05 | 0.0000 | 18 | -1.76 | 0.0000 | 26 | -1.90 | 0.0000 | 148 | -2.44 | 0.0000 |
| REACTOME_EXTENSION_OF_TELOMERES | 37 | -2.04 | 0.0000 | 89 | -1.69 | 0.0001 | 33 | -1.89 | 0.0000 | 69 | -2.62 | 0.0000 |
| KEGG_DNA_REPLICATION | 31 | -2.05 | 0.0000 | 28 | -1.76 | 0.0000 | 34 | -1.89 | 0.0000 | 137 | -2.46 | 0.0000 |
| REACTOME_HDR_THROUGH_HOMOLOGOUS_RECOMBINATION_HRR | 35 | -2.05 | 0.0000 | 88 | -1.69 | 0.0001 | 58 | -1.84 | 0.0000 | 53 | -2.67 | 0.0000 |
| HALLMARK_MYC_TARGETS_V2 | 36 | -2.05 | 0.0000 | 40 | -1.73 | 0.0000 | 125 | -1.80 | 0.0000 | 44 | -2.74 | 0.0000 |
| REACTOME_FORMATION_OF_RNA_POL_II_ELONGATION_COMPLEX | 30 | -2.05 | 0.0000 | 44 | -1.73 | 0.0000 | 57 | -1.85 | 0.0000 | 115 | -2.49 | 0.0000 |
| REACTOME_RNA_POLYMERASE_II_TRANSCRIBES_SNRNA_GENES | 86 | -1.97 | 0.0000 | 59 | -1.71 | 0.0000 | 85 | -1.82 | 0.0000 | 39 | -2.78 | 0.0000 |
| REACTOME_ORC1_REMOVAL_FROM_CHROMATIN | 82 | -1.98 | 0.0000 | 71 | -1.70 | 0.0001 | 47 | -1.85 | 0.0000 | 70 | -2.62 | 0.0000 |
| REACTOME_TRANSPORT_OF_MATURE_TRANSCRIPT_TO_CYTOPLASM | 81 | -1.98 | 0.0000 | 80 | -1.70 | 0.0001 | 81 | -1.82 | 0.0000 | 32 | -2.81 | 0.0000 |
| WONG_EMBRYONIC_STEM_CELL_CORE | 61 | -2.00 | 0.0000 | 127 | -1.67 | 0.0002 | 64 | -1.84 | 0.0000 | 27 | -2.86 | 0.0000 |
| REACTOME_NUCLEOTIDE_EXCISION_REPAIR | 45 | -2.02 | 0.0000 | 92 | -1.69 | 0.0001 | 56 | -1.85 | 0.0000 | 90 | -2.55 | 0.0000 |
| REACTOME_HIV_LIFE_CYCLE | 55 | -2.00 | 0.0000 | 110 | -1.69 | 0.0001 | 78 | -1.83 | 0.0000 | 47 | -2.72 | 0.0000 |
| WP_TRANSLATION_FACTORS | 125 | -1.93 | 0.0000 | 22 | -1.76 | 0.0000 | 53 | -1.85 | 0.0000 | 96 | -2.54 | 0.0000 |

| | K562 | | | ST486 | | | HepG2 | | | MCF7 | | |
|---|---|---|---|---|---|---|---|---|---|---|---|---|
| **GENE SETS** | **RANK** | **NES** | **FDR q-val** | **RANK** | **NES** | **FDR q-val** | **RANK** | **NES** | **FDR q-val** | **RANK** | **NES** | **FDR q-val** |
| DANG_MYC_TARGETS_UP | 40 | -2.04 | 0.0000 | 161 | -1.66 | 0.0003 | 62 | -1.84 | 0.0000 | 34 | -2.79 | 0.0000 |
| MANALO_HYPOXIA_DN | 59 | -2.00 | 0.0000 | 138 | -1.67 | 0.0002 | 82 | -1.82 | 0.0000 | 26 | -2.86 | 0.0000 |
| REACTOME_S_PHASE | 73 | -1.98 | 0.0000 | 105 | -1.69 | 0.0001 | 93 | -1.82 | 0.0000 | 37 | -2.78 | 0.0000 |
| REACTOME_TRANSCRIPTION_OF_THE_HIV_GENOME | 33 | -2.05 | 0.0000 | 45 | -1.73 | 0.0000 | 69 | -1.83 | 0.0000 | 162 | -2.41 | 0.0000 |
| REACTOME_HIV_INFECTION | 98 | -1.95 | 0.0000 | 82 | -1.70 | 0.0001 | 112 | -1.80 | 0.0000 | 33 | -2.81 | 0.0000 |
| REACTOME_FORMATION_OF_TC_NER_PRE_INCISION_COMPLEX | 72 | -1.98 | 0.0000 | 43 | -1.73 | 0.0000 | 25 | -1.91 | 0.0000 | 190 | -2.38 | 0.0000 |
| REACTOME_HOMOLOGY_DIRECTED_REPAIR | 50 | -2.01 | 0.0000 | 166 | -1.66 | 0.0003 | 95 | -1.81 | 0.0000 | 40 | -2.77 | 0.0000 |
| REACTOME_G2_M_CHECKPOINTS | 75 | -1.98 | 0.0000 | 128 | -1.67 | 0.0002 | 127 | -1.79 | 0.0000 | 29 | -2.85 | 0.0000 |
| REACTOME_TELOMERE_C_STRAND_LAGGING_STRAND_SYNTHESIS | 70 | -1.99 | 0.0000 | 108 | -1.69 | 0.0001 | 38 | -1.89 | 0.0000 | 144 | -2.45 | 0.0000 |
| REACTOME_ACTIVATION_OF_ATR_IN_RESPONSE_TO_REPLICATION_STRESS | 47 | -2.02 | 0.0000 | 106 | -1.69 | 0.0001 | 80 | -1.83 | 0.0000 | 132 | -2.46 | 0.0000 |
| REACTOME_CELL_CYCLE_CHECKPOINTS | 106 | -1.94 | 0.0000 | 126 | -1.67 | 0.0002 | 118 | -1.80 | 0.0000 | 16 | -2.97 | 0.0000 |
| REACTOME_FORMATION_OF_THE_EARLY_ELONGATION_COMPLEX | 56 | -2.00 | 0.0000 | 19 | -1.76 | 0.0000 | 30 | -1.90 | 0.0000 | 269 | -2.25 | 0.0000 |
| IRITANI_MAD1_TARGETS_DN | 89 | -1.96 | 0.0000 | 38 | -1.74 | 0.0000 | 128 | -1.79 | 0.0000 | 125 | -2.47 | 0.0000 |
| WP_EUKARYOTIC_TRANSCRIPTION_INITIATION | 51 | -2.01 | 0.0000 | 34 | -1.74 | 0.0000 | 84 | -1.82 | 0.0000 | 216 | -2.33 | 0.0000 |
| REN_BOUND_BY_E2F | 32 | -2.05 | 0.0000 | 184 | -1.64 | 0.0004 | 50 | -1.85 | 0.0000 | 135 | -2.46 | 0.0000 |
| CROONQUIST_NRAS_SIGNALING_DN | 48 | -2.02 | 0.0000 | 197 | -1.64 | 0.0006 | 65 | -1.84 | 0.0000 | 113 | -2.50 | 0.0000 |
| REACTOME_TRANSPORT_OF_MATURE_MRNAS_DERIVED_FROM_INTRONLESS_TRANSCRIPTS | 111 | -1.94 | 0.0000 | 70 | -1.70 | 0.0001 | 207 | -1.75 | 0.0001 | 48 | -2.70 | 0.0000 |
| KEGG_RNA_POLYMERASE | 84 | -1.97 | 0.0000 | 129 | -1.67 | 0.0002 | 37 | -1.89 | 0.0000 | 203 | -2.35 | 0.0000 |
| REACTOME_SEPARATION_OF_SISTER_CHROMATIDS | 196 | -1.88 | 0.0000 | 104 | -1.69 | 0.0001 | 147 | -1.78 | 0.0000 | 28 | -2.85 | 0.0000 |
| PID_ATR_PATHWAY | 179 | -1.89 | 0.0000 | 49 | -1.72 | 0.0000 | 120 | -1.80 | 0.0000 | 140 | -2.45 | 0.0000 |
| MODY_HIPPOCAMPUS_PRENATAL | 44 | -2.02 | 0.0000 | 250 | -1.62 | 0.001 | 132 | -1.79 | 0.0000 | 75 | -2.61 | 0.0000 |
| REACTOME_DNA_DOUBLE_STRAND_BREAK_REPAIR | 68 | -1.99 | 0.0000 | 201 | -1.64 | 0.0006 | 185 | -1.76 | 0.0001 | 49 | -2.70 | 0.0000 |
| REACTOME_PROCESSING_OF_CAPPED_INTRONLESS_PRE_MRNA | 49 | -2.01 | 0.0000 | 155 | -1.66 | 0.0003 | 71 | -1.83 | 0.0000 | 243 | -2.29 | 0.0000 |
| HALLMARK_E2F_TARGETS | 93 | -1.96 | 0.0000 | 217 | -1.63 | 0.0007 | 172 | -1.77 | 0.0001 | 45 | -2.74 | 0.0000 |
| REACTOME_HOST_INTERACTIONS_OF_HIV_FACTORS | 200 | -1.88 | 0.0000 | 121 | -1.68 | 0.0001 | 171 | -1.77 | 0.0001 | 38 | -2.78 | 0.0000 |
| REACTOME_POSTMITOTIC_NUCLEAR_PORE_COMPLEX_NPC_REFORMATION | 228 | -1.86 | 0.0000 | 36 | -1.74 | 0.0000 | 176 | -1.77 | 0.0001 | 104 | -2.52 | 0.0000 |
| REACTOME_MITOCHONDRIAL_TRNA_AMINOACYLATION | 137 | -1.92 | 0.0000 | 32 | -1.74 | 0.0000 | 102 | -1.81 | 0.0000 | 292 | -2.22 | 0.0000 |
| REACTOME_MITOTIC_METAPHASE_AND_ANAPHASE | 233 | -1.86 | 0.0000 | 151 | -1.66 | 0.0002 | 156 | -1.78 | 0.0000 | 24 | -2.86 | 0.0000 |
| REACTOME_TRNA_AMINOACYLATION | 104 | -1.95 | 0.0000 | 29 | -1.75 | 0.0000 | 157 | -1.78 | 0.0000 | 291 | -2.22 | 0.0000 |
| REACTOME_MITOTIC_SPINDLE_CHECKPOINT | 220 | -1.87 | 0.0000 | 143 | -1.67 | 0.0002 | 184 | -1.76 | 0.0001 | 41 | -2.74 | 0.0000 |
| KEGG_AMINOACYL_TRNA_BIOSYNTHESIS | 69 | -1.99 | 0.0000 | 33 | -1.74 | 0.0000 | 224 | -1.74 | 0.0002 | 263 | -2.27 | 0.0000 |
| REACTOME_CELL_CYCLE_MITOTIC | 198 | -1.88 | 0.0000 | 189 | -1.64 | 0.0005 | 200 | -1.75 | 0.0001 | 43 | -2.74 | 0.0000 |
| REACTOME_M_PHASE | 254 | -1.85 | 0.0000 | 190 | -1.64 | 0.0005 | 219 | -1.74 | 0.0001 | 50 | -2.70 | 0.0000 |
| REACTOME_REGULATION_OF_TP53_ACTIVITY_THROUGH_PHOSPHORYLATION | 43 | -2.02 | 0.0000 | 171 | -1.65 | 0.0003 | 309 | -1.70 | 0.0006 | 198 | -2.36 | 0.0000 |
| KEGG_OXIDATIVE_PHOSPHORYLATION | 150 | -1.91 | 0.0000 | 367 | -1.58 | 0.003 | 42 | -1.87 | 0.0000 | 255 | -2.28 | 0.0000 |
| REACTOME_RESPIRATORY_ELECTRON_TRANSPORT | 114 | -1.93 | 0.0000 | 561 | -1.51 | 0.0146 | 45 | -1.86 | 0.0000 | 122 | -2.48 | 0.0000 |
| REACTOME_RNA_POLYMERASE_I_TRANSCRIPTION_TERMINATION | 103 | -1.95 | 0.0000 | 46 | -1.73 | 0.0000 | 76 | -1.83 | 0.0000 | 640 | -1.84 | 0.0052 |
| XU_RESPONSE_TO_TRETINOIN_AND_NSC682994_DN | 299 | -1.82 | 0.0001 | 25 | -1.76 | 0.0000 | 202 | -1.75 | 0.0001 | 342 | -2.17 | 0.0000 |
| REACTOME_MICRORNA_MIRNA_BIOGENESIS | 408 | -1.75 | 0.0007 | 160 | -1.66 | 0.0003 | 46 | -1.85 | 0.0000 | 326 | -2.19 | 0.0000 |
| MOOTHA_VOXPHOS | 262 | -1.84 | 0.0001 | 493 | -1.54 | 0.0082 | 44 | -1.86 | 0.0000 | 217 | -2.32 | 0.0000 |
| WP_ELECTRON_TRANSPORT_CHAIN_OXPHOS_SYSTEM_IN_MITOCHONDRIA | 232 | -1.86 | 0.0000 | 505 | -1.53 | 0.0092 | 43 | -1.87 | 0.0000 | 395 | -2.11 | 0.0001 |
| REACTOME_NUCLEOBASE_BIOSYNTHESIS | 453 | -1.73 | 0.0012 | 50 | -1.72 | 0.0000 | 528 | -1.60 | 0.0072 | 654 | -1.82 | 0.0065 |
| WP_MITOCHONDRIAL_COMPLEX_I_ASSEMBLY_MODEL_OXPHOS_SYSTEM | 263 | -1.84 | 0.0001 | 1359 | -1.32 | 0.1732 | 49 | -1.85 | 0.0000 | 293 | -2.22 | 0.0000 |
| WANG_ADIPOGENIC_GENES_REPRESSED_BY_SIRT1 | 629 | -1.65 | 0.0067 | 48 | -1.72 | 0.0000 | 814 | -1.50 | 0.0369 | 792 | -1.71 | 0.0211 |

**Table S7.** Top 50 enriched gene sets among genes adjacent to essential E-boxes.

| | K562 | | | ST486 | | | HepG2 | | | MCF7 | | |
|---|---|---|---|---|---|---|---|---|---|---|---|---|
| PROCESSES | RANK | NES | FDR q-val | RANK | NES | FDR q-val | RANK | NES | FDR q-val | RANK | NES | FDR q-val |
| REACTOME_TRANSLATION | 2 | -1,72 | 0.003 | 19 | -1,41 | 0.2655 | 17 | -1,54 | 0.0324 | 3 | -2,12 | 0.000 |
| REACTOME_METABOLISM_OF_RNA | 6 | -1,67 | 0.0065 | 8 | -1,45 | 0.2156 | 19 | -1,52 | 0.0451 | 2 | -2,24 | 0.000 |
| REACTOME_RRNA_PROCESSING | 3 | -1,71 | 0.002 | 35 | -1,37 | 0.2988 | 18 | -1,53 | 0.0372 | 1 | -2,25 | 0.000 |
| REACTOME_SIGNALING_BY_ROBO_RECEPTORS | 13 | -1,64 | 0.0084 | 34 | -1,37 | 0.307 | 6 | -1,57 | 0.021 | 6 | -2,04 | 0.000 |
| REACTOME_INFLUENZA_INFECTION | 8 | -1,66 | 0.0086 | 31 | -1,37 | 0.3028 | 16 | -1,55 | 0.0212 | 11 | -1,95 | 0.0012 |
| HSIAO_HOUSEKEEPING_GENES | 27 | -1,58 | 0.034 | 17 | -1,42 | 0.2092 | 12 | -1,55 | 0.0238 | 10 | -1,95 | 0.0012 |
| REACTOME_REGULATION_OF_EXPRESSION_OF_SLITS_AND_ROBOS | 21 | -1,60 | 0.0205 | 37 | -1,37 | 0.2875 | 2 | -1,59 | 0.0285 | 7 | -2,00 | 0.0005 |
| REACTOME_CELLULAR_RESPONSE_TO_STARVATION | 11 | -1,64 | 0.0096 | 39 | -1,37 | 0.2793 | 1 | -1,59 | 0.0551 | 23 | -1,88 | 0.0032 |
| REACTOME_EUKARYOTIC_TRANSLATION_INITIATION | 19 | -1,61 | 0.0177 | 42 | -1,36 | 0.2629 | 9 | -1,56 | 0.0229 | 4 | -2,09 | 0.000 |
| WP_CYTOPLASMIC_RIBOSOMAL_PROTEINS | 5 | -1,67 | 0.0078 | 52 | -1,34 | 0.3208 | 3 | -1,59 | 0.0241 | 17 | -1,92 | 0.002 |
| KEGG_RIBOSOME | 12 | -1,64 | 0.0091 | 49 | -1,34 | 0.3305 | 10 | -1,56 | 0.0209 | 9 | -1,96 | 0.0012 |
| REACTOME_RESPONSE_OF_EIF2AK4_GCN2_TO_AMINO_ACID_DEFICIENCY | 15 | -1,64 | 0.0077 | 50 | -1,34 | 0.327 | 4 | -1,58 | 0.0199 | 13 | -1,94 | 0.0014 |
| REACTOME_EUKARYOTIC_TRANSLATION_ELONGATION | 18 | -1,62 | 0.0129 | 45 | -1,35 | 0.322 | 7 | -1,57 | 0.0192 | 12 | -1,95 | 0.0014 |
| REACTOME_NONSENSE_MEDIATED_DECAY_NMD | 22 | -1,60 | 0.0196 | 43 | -1,36 | 0.2843 | 13 | -1,55 | 0.025 | 5 | -2,07 | 0.000 |
| CAIRO_HEPATOBLASTOMA_CLASSES_UP | 10 | -1,65 | 0.0092 | 11 | -1,44 | 0.1928 | 45 | -1,41 | 0.2831 | 20 | -1,90 | 0.0025 |
| REACTOME_SELENOAMINO_ACID_METABOLISM | 4 | -1,68 | 0.0091 | 59 | -1,33 | 0.3106 | 14 | -1,55 | 0.0237 | 18 | -1,92 | 0.0021 |
| BERENJENO_TRANSFORMED_BY_RHOA_UP | 24 | -1,60 | 0.0203 | 15 | -1,43 | 0.2146 | 36 | -1,43 | 0.2413 | 21 | -1,89 | 0.0028 |
| WANG_TUMOR_INVASIVENESS_UP | 26 | -1,58 | 0.0303 | 7 | -1,45 | 0.2385 | 29 | -1,44 | 0.2351 | 34 | -1,79 | 0.0174 |
| REACTOME_SRP_DEPENDENT_COTRANSLATIONAL_PROTEIN_TARGETING_TO_MEMBRANE | 14 | -1,64 | 0.0083 | 48 | -1,34 | 0.3371 | 20 | -1,52 | 0.0492 | 15 | -1,94 | 0.0014 |
| DODD_NASOPHARYNGEAL_CARCINOMA_DN | 30 | -1,56 | 0.0571 | 9 | -1,45 | 0.1996 | 39 | -1,41 | 0.2917 | 28 | -1,84 | 0.0073 |
| REACTOME_METABOLISM_OF_AMINO_ACIDS_AND_DERIVATIVES | 17 | -1,63 | 0.0114 | 41 | -1,36 | 0.2686 | 8 | -1,57 | 0.0212 | 45 | -1,71 | 0.0587 |
| LEE_BMP2_TARGETS_DN | 9 | -1,66 | 0.0081 | 12 | -1,44 | 0.214 | 69 | -1,35 | 0.4071 | 24 | -1,85 | 0.0063 |
| HALLMARK_MYC_TARGETS_V1 | 28 | -1,57 | 0.0424 | 3 | -1,50 | 0.114 | 58 | -1,38 | 0.3373 | 41 | -1,72 | 0.0506 |
| REACTOME_NERVOUS_SYSTEM_DEVELOPMENT | 49 | -1,50 | 0.1514 | 68 | -1,32 | 0.3286 | 5 | -1,58 | 0.0222 | 14 | -1,94 | 0.0014 |
| WONG_EMBRYONIC_STEM_CELL_CORE | 7 | -1,66 | 0.0078 | 98 | -1,29 | 0.3406 | 25 | -1,45 | 0.2048 | 8 | -1,96 | 0.0014 |
| REACTOME_INFECTIOUS_DISEASE | 16 | -1,63 | 0.0105 | 53 | -1,34 | 0.319 | 11 | -1,56 | 0.0224 | 61 | 0,00 | 0.000 |
| REACTOME_CELLULAR_RESPONSES_TO_EXTERNAL_STIMULI | 31 | -1,56 | 0.0561 | 58 | -1,33 | 0.313 | 15 | -1,55 | 0.0223 | 49 | -1,70 | 0.0677 |
| PUJANA_BRCA1_PCC_NETWORK | 34 | -1,55 | 0.0564 | 27 | -1,38 | 0.2877 | 54 | -1,39 | 0.2867 | 57 | 0,00 | 0.000 |
| TIEN_INTESTINE_PROBIOTICS_24HR_DN | 38 | -1,51 | 0.1435 | 94 | -1,29 | 0.3332 | 21 | -1,51 | 0.0656 | 90 | 0,00 | 0.000 |
| PUJANA_CHEK2_PCC_NETWORK | 32 | -1,55 | 0.0575 | 115 | -1,27 | 0.3674 | 70 | -1,35 | 0.4194 | 31 | -1,82 | 0.0113 |
| LI_AMPLIFIED_IN_LUNG_CANCER | 127 | -1,39 | 0.3328 | 23 | -1,39 | 0.3066 | 47 | -1,40 | 0.2771 | 62 | 0,00 | 0.000 |
| JISON_SICKLE_CELL_DISEASE_DN | 46 | -1,51 | 0.1329 | 174 | -1,21 | 0.4354 | 31 | -1,44 | 0.2311 | 16 | -1,94 | 0.0014 |
| REACTOME_DEVELOPMENTAL_BIOLOGY | 117 | -1,40 | 0.3227 | 70 | -1,32 | 0.3318 | 46 | -1,40 | 0.2797 | 38 | -1,75 | 0.0362 |
| OSMAN_BLADDER_CANCER_DN | 1 | -1,72 | 0.006 | 108 | -1,27 | 0.3717 | 122 | -1,28 | 0.5499 | 42 | -1,72 | 0.0509 |
| FISCHER_DREAM_TARGETS | 39 | -1,51 | 0.1448 | 56 | -1,33 | 0.3169 | 171 | -1,24 | 0.5691 | 26 | -1,84 | 0.0077 |
| DUTERTRE_ESTRADIOL_RESPONSE_6HR_UP | 132 | -1,39 | 0.3351 | 75 | -1,31 | 0.3322 | 35 | -1,43 | 0.2319 | 58 | 0,00 | 0.000 |
| MANALO_HYPOXIA_DN | 86 | -1,44 | 0.2568 | 102 | -1,28 | 0.3462 | 118 | -1,28 | 0.553 | 19 | -1,92 | 0.0021 |
| KARLSSON_TGFB1_TARGETS_UP | 111 | -1,41 | 0.3038 | 107 | -1,27 | 0.365 | 104 | -1,29 | 0.5435 | 22 | -1,89 | 0.0028 |
| GARY_CD5_TARGETS_DN | 79 | -1,45 | 0.2469 | 202 | -1,19 | 0.4502 | 38 | -1,42 | 0.2825 | 30 | -1,82 | 0.0104 |
| KINSEY_TARGETS_OF_EWSR1_FLII_FUSION_UP | 81 | -1,44 | 0.2586 | 29 | -1,38 | 0.3112 | 232 | -1,19 | 0.6121 | 40 | -1,73 | 0.0507 |
| RODRIGUES_THYROID_CARCINOMA_POORLY_DIFFERENTIATED_UP | 66 | -1,46 | 0.2218 | 104 | -1,28 | 0.3575 | 48 | -1,40 | 0.2714 | 227 | 0,00 | 0.000 |
| FOURNIER_ACINAR_DEVELOPMENT_LATE_2 | 183 | -1,34 | 0.4391 | 21 | -1,39 | 0.3041 | 128 | -1,27 | 0.5785 | 119 | 0,00 | 0.000 |
| MONNIER_POSTRADIATION_TUMOR_ESCAPE_UP | 35 | -1,54 | 0.0741 | 135 | -1,24 | 0.4069 | 215 | -1,20 | 0.6066 | 74 | 0,00 | 0.000 |
| SENGUPTA_NASOPHARYNGEAL_CARCINOMA_UP | 230 | -1,30 | 0.5218 | 14 | -1,43 | 0.2069 | 75 | -1,34 | 0.4164 | 176 | 0,00 | 0.000 |
| MARTENS_TRETINOIN_RESPONSE_DN | 25 | -1,59 | 0.025 | 238 | -1,15 | 0.5105 | 195 | -1,22 | 0.5705 | 43 | -1,72 | 0.053 |
| KIM_ALL_DISORDERS_OLIGODENDROCYTE_NUMBER_CORR_UP | 47 | -1,50 | 0.1457 | 22 | -1,39 | 0.2978 | 63 | -1,37 | 0.3542 | 371 | 0,00 | 0.000 |
| REACTOME_PROCESSING_OF_CAPPED_INTRON_CONTAINING_PRE_MRNA | 283 | -1,24 | 0.654 | 110 | -1,27 | 0.3682 | 78 | -1,34 | 0.4316 | 32 | -1,80 | 0.0162 |
| MOOTHA_HUMAN_MITODB_6_2002 | 41 | -1,51 | 0.1413 | 118 | -1,26 | 0.3832 | 220 | -1,19 | 0.6066 | 125 | 0,00 | 0.000 |
| KEGG_PURINE_METABOLISM | 107 | -1,41 | 0.3088 | 66 | -1,33 | 0.3064 | 290 | -1,13 | 0.7009 | 44 | -1,72 | 0.0531 |
| REACTOME_MRNA_SPLICING | 351 | -1,18 | 0.742 | 186 | -1,20 | 0.437 | 34 | -1,43 | 0.2328 | 36 | -1,77 | 0.0248 |
| YAGI_AML_WITH_INV_16_TRANSLOCATION | 29 | -1,56 | 0.0506 | 164 | -1,22 | 0.4216 | 196 | -1,22 | 0.5746 | 266 | 0,00 | 0.000 |
| MUELLER_PLURINET | 56 | -1,48 | 0.1869 | 362 | -0,96 | 0.8146 | 209 | -1,20 | 0.5963 | 33 | -1,79 | 0.0176 |
| DESERT_STEM_CELL_HEPATOCELLULAR_CARCINOMA_SUBCLASS_UP | 136 | -1,39 | 0.3375 | 16 | -1,43 | 0.2056 | 126 | -1,27 | 0.5787 | 418 | 0,00 | 0.000 |
| REACTOME_CHROMATIN_MODIFYING_ENZYMES | 33 | -1,55 | 0.0563 | 341 | -1,01 | 0.7537 | 149 | -1,26 | 0.5413 | 211 | 0,00 | 0.000 |
| BASAKI_YBX1_TARGETS_UP | 238 | -1,28 | 0.5525 | 13 | -1,43 | 0.2172 | 168 | -1,24 | 0.5479 | 332 | 0,00 | 0.000 |

| | K562 | | | ST486 | | | HepG2 | | | MCF7 | | |
|---|---|---|---|---|---|---|---|---|---|---|---|---|
| PROCESSES | RANK | NES | FDR q-val | RANK | NES | FDR q-val | RANK | NES | FDR q-val | RANK | NES | FDR q-val |
| IVANOVA_HEMATOPOIESIS_EARLY_PROGENITOR | 45 | -1,51 | 0.1329 | 103 | -1,28 | 0.3531 | 210 | -1,20 | 0.5972 | 394 | 0,00 | 0.000 |
| GRYDER_PAX3FOXO1_TOP_ENHANCERS | 385 | -1,16 | 0.7788 | 4 | -1,47 | 0.2276 | 270 | -1,14 | 0.6886 | 100 | 0,00 | 0.000 |
| LASTOWSKA_NEUROBLASTOMA_COPY_NUMBER_UP | 42 | -1,51 | 0.1412 | 145 | -1,24 | 0.4113 | 272 | -1,14 | 0.6907 | 300 | 0,00 | 0.000 |
| BENPORATH_MYC_MAX_TARGETS | 503 | -1,04 | 0.926 | 20 | -1,40 | 0.2568 | 108 | -1,29 | 0.5543 | 197 | 0,00 | 0.000 |
| WANG_RESPONSE_TO_GSK3_INHIBITOR_SB216763_DN | 370 | -1,17 | 0.7582 | 44 | -1,35 | 0.2915 | 83 | -1,33 | 0.4607 | 347 | 0,00 | 0.000 |
| MARTINEZ_RB1_AND_TP53_TARGETS_DN | 355 | -1,18 | 0.7414 | 36 | -1,37 | 0.2946 | 217 | -1,20 | 0.6084 | 254 | 0,00 | 0.000 |
| RHEIN_ALL_GLUCOCORTICOID_THERAPY_DN | 23 | -1,60 | 0.0193 | 156 | -1,23 | 0.4058 | 94 | -1,31 | 0.4875 | 606 | 0,00 | 0.000 |
| KIM_BIPOLAR_DISORDER_OLIGODENDROCYTE_DENSITY_CORR_UP | 65 | -1,46 | 0.2154 | 133 | -1,25 | 0.405 | 23 | -1,46 | 0.2064 | 708 | 0,00 | 0.000 |
| BLALOCK_ALZHEIMERS_DISEASE_DN | 167 | -1,36 | 0.3934 | 71 | -1,31 | 0.3299 | 28 | -1,44 | 0.2328 | 691 | 0,00 | 0.000 |
| TARTE_PLASMA_CELL_VS_PLASMABLAST_UP | 161 | -1,36 | 0.3886 | 28 | -1,38 | 0.2841 | 457 | -0,95 | 0.8906 | 319 | 0,00 | 0.000 |
| BUYTAERT_PHOTODYNAMIC_THERAPY_STRESS_DN | 431 | -1,12 | 0.8382 | 40 | -1,36 | 0.2746 | 164 | -1,24 | 0.5449 | 338 | 0,00 | 0.000 |
| STARK_PREFRONTAL_CORTEX_22Q11_DELETION_DN | 20 | -1,61 | 0.0178 | 158 | -1,23 | 0.41 | 43 | -1,41 | 0.2884 | 790 | 0,00 | 0.000 |
| MARTINEZ_TP53_TARGETS_DN | 334 | -1,19 | 0.7369 | 10 | -1,44 | 0.2066 | 251 | -1,16 | 0.6544 | 452 | 0,00 | 0.000 |
| GRYDER_PAX3FOXO1_ENHANCERS_IN_TADS | 569 | -0,99 | 0.9607 | 38 | -1,37 | 0.2822 | 346 | -1,06 | 0.7977 | 123 | 0,00 | 0.000 |
| REACTOME_CYTOKINE_SIGNALING_IN_IMMUNE_SYSTEM | 321 | -1,21 | 0.7069 | 33 | -1,37 | 0.3046 | 487 | -0,89 | 0.9443 | 419 | 0,00 | 0.000 |
| BOUDOUKHA_BOUND_BY_IGF2BP2 | 43 | -1,51 | 0.1384 | 219 | -1,17 | 0.4797 | 123 | -1,27 | 0.5558 | 1049 | 0,00 | 0.000 |
| BAELDE_DIABETIC_NEPHROPATHY_DN | 489 | -1,05 | 0.9311 | 245 | -1,15 | 0.5166 | 40 | -1,41 | 0.2965 | 677 | 0,00 | 0.000 |
| PHONG_TNF_RESPONSE_VIA_P38_COMPLETE | 97 | -1,43 | 0.2725 | 46 | -1,35 | 0.3159 | 288 | -1,13 | 0.7026 | 1148 | 0,00 | 0.000 |
| LIU_SOX4_TARGETS_DN | 556 | -1,01 | 0.9379 | 5 | -1,47 | 0.2098 | 389 | -1,01 | 0.8655 | 712 | 0,00 | 0.000 |
| ACEVEDO_NORMAL_TISSUE_ADJACENT_TO_LIVER_TUMOR_DN | 513 | -1,03 | 0.9373 | 18 | -1,41 | 0.2512 | 205 | -1,21 | 0.5789 | 1036 | 0,00 | 0.000 |
| BOQUEST_STEM_CELL_CULTURED_VS_FRESH_UP | 395 | -1,15 | 0.7783 | 26 | -1,38 | 0.2956 | 462 | -0,94 | 0.893 | 1082 | 0,00 | 0.000 |
| REACTOME_SIGNALING_BY_WNT | 673 | -0,87 | 1.0000 | 467 | -0,59 | 1.0000 | 41 | -1,41 | 0.299 | 1032 | 0,00 | 0.000 |
| BLANCO_MELO_INFLUENZA_A_INFECTION_A594_CELLS_DN | 780 | -0,77 | 1.0000 | 436 | -0,75 | 0.9705 | 30 | -1,44 | 0.2306 | 1165 | 0,00 | 0.000 |
| ALONSO_METASTASIS_UP | | | | 348 | -0,99 | 0.779 | | | | 35 | -1,78 | 0.0201 |
| BILANGES_SERUM_AND_RAPAMYCIN_SENSITIVE_GENES | | | | | | | 22 | -1,46 | 0.2011 | 50 | -1,70 | 0.0673 |
| BOYAULT_LIVER_CANCER_SUBCLASS_G3_UP | 75 | -1,45 | 0.248 | | | | 26 | -1,45 | 0.2072 | 498 | 0,00 | 0.000 |
| BRUECKNER_TARGETS_OF_MIRLET7A3_UP | | | | | | | | | | 39 | -1,73 | 0.045 |
| CHICAS_RB1_TARGETS_GROWING | 226 | -1,30 | 0.5105 | | | | 24 | -1,46 | 0.2031 | 85 | 0,00 | 0.000 |
| DANG_MYC_TARGETS_UP | 141 | -1,38 | 0.361 | 24 | -1,39 | 0.3107 | | | | 47 | -1,71 | 0.0607 |
| GAVIN_FOXP3_TARGETS_CLUSTER_T7 | 36 | -1,54 | 0.082 | | | | | | | 95 | 0,00 | 0.000 |
| GRAESSMANN_APOPTOSIS_BY_SERUM_DEPRIVATION_UP | 723 | -0,83 | 1.0000 | | | | 37 | -1,43 | 0.244 | 236 | 0,00 | 0.000 |
| HOLLEMAN_VINCRISTINE_RESISTANCE_B_ALL_DN | | | | 30 | -1,37 | 0.3098 | | | | 305 | 0,00 | 0.000 |
| HOLLMANN_APOPTOSIS_VIA_CD40_DN | 592 | -0,97 | 0.967 | | | | 50 | -1,40 | 0.289 | 694 | 0,00 | 0.000 |
| LEE_EARLY_T_LYMPHOCYTE_UP | 40 | -1,51 | 0.1446 | | | | | | | 896 | 0,00 | 0.000 |
| MALONEY_RESPONSE_TO_17AAG_DN | 346 | -1,19 | 0.7482 | 47 | -1,34 | 0.3323 | | | | 536 | 0,00 | 0.000 |
| MARTINEZ_RESPONSE_TO_TRABECTEDIN_DN | 119 | -1,40 | 0.3242 | | | | | | | 27 | -1,84 | 0.0075 |
| MARZEC_IL2_SIGNALING_UP | 37 | -1,54 | 0.0817 | | | | | | | 543 | 0,00 | 0.000 |
| NIKOLSKY_BREAST_CANCER_8Q23_Q24_AMPLICON | | | | 389 | -0,91 | 0.8701 | 49 | -1,40 | 0.277 | 1364 | 0,00 | 0.000 |
| REACTOME_ACTIVATION_OF_THE_MRNA_UPON_BINDING_OF_THE_CAP_BINDING_COMPLEX_AND_EIFS_AND_SUBSEQUENT_BINDING_TO_43S | | | | | | | | | | 37 | -1,76 | 0.0317 |
| REACTOME_ASPARAGINE_N_LINKED_GLYCOSYLATION | 609 | -0,95 | 0.976 | | | | 27 | -1,45 | 0.2117 | 823 | 0,00 | 0.000 |
| REACTOME_DISEASES_OF_GLYCOSYLATION | 48 | -1,50 | 0.1504 | 1 | -1,52 | 0.1389 | | | | 924 | 0,00 | 0.000 |
| REACTOME_DISEASES_OF_METABOLISM | 126 | -1,39 | 0.3324 | 2 | -1,51 | 0.1202 | | | | 577 | 0,00 | 0.000 |
| REACTOME_HIV_INFECTION | 206 | -1,32 | 0.4834 | | | | 53 | -1,39 | 0.2903 | 46 | -1,71 | 0.0585 |
| REACTOME_INTERFERON_SIGNALING | 449 | -1,09 | 0.8777 | 32 | -1,37 | 0.3095 | | | | 439 | 0,00 | 0.000 |
| REACTOME_MITOCHONDRIAL_TRANSLATION | 98 | -1,43 | 0.27 | | | | | | | 48 | -1,70 | 0.063 |
| REACTOME_RRNA_MODIFICATION_IN_THE_NUCLEUS_AND_CYTOSOL | 95 | -1,43 | 0.2705 | | | | 194 | -1,22 | 0.5636 | 29 | -1,83 | 0.0081 |
| REACTOME_SIGNALING_BY_RECEPTOR_TYROSINE_KINASES | 696 | -0,85 | 1.0000 | 6 | -1,46 | 0.2421 | | | | 928 | 0,00 | 0.000 |
| REACTOME_TCF_DEPENDENT_SIGNALING_IN_RESPONSE_TO_WNT | 638 | -0,91 | 1.0000 | | | | 44 | -1,41 | 0.2828 | 457 | 0,00 | 0.000 |
| SCHLOSSER_SERUM_RESPONSE_DN | 433 | -1,11 | 0.8412 | | | | 42 | -1,41 | 0.2949 | 837 | 0,00 | 0.000 |
| TIEN_INTESTINE_PROBIOTICS_6HR_UP | 44 | -1,51 | 0.1353 | | | | | | | 25 | -1,85 | 0.0072 |
| WANG_PROSTATE_CANCER_ANDROGEN_INDEPENDENT | | | | 25 | -1,39 | 0.3019 | | | | 261 | 0,00 | 0.000 |
| WP_16P112_PROXIMAL_DELETION_SYNDROME | 50 | -1,50 | 0.1502 | | | | | | | 709 | 0,00 | 0.000 |
| WP_PATHWAYS_AFFECTED_IN_ADENOID_CYSTIC_CARCINOMA | | | | | | | 32 | -1,44 | 0.2343 | | | |
| YAGI_AML_SURVIVAL | | | | 265 | -1,13 | 0.5477 | 33 | -1,43 | 0.238 | 654 | 0,00 | 0.000 |

**Table S8.** E-box editing in K562 cells with CRISPR/Cas9: +1 insertions.

| sgRNA | E-box | % +1 insertions | % +G/C | % E-box not changed | Most prevalent E-box after +1 insertion |
|---|---|---|---|---|---|
| **Chr17_BS377_sg1** | CACGTG | 9 | 20 | 1,8 | CAACGTG |
| **Chr17_BS377_sg2** | CGCGTG | 65 | 78 | 50 | CGCGTGG |
| **Chr17_BS377_sg3** | CGCGTG | 45 | 0 | 0 (cut in the middle) | - |
| **Chr11_BS2113_sg1** | CACATG | 60 | 80 | 48 | CACATGG |
| **Chr10_BS212_sg1** | CACATG | 80 | 70 | 56 | CACATGG |
| **Chr3_BS897_sg1** | CATGTG | 0 | 0 | 0 | - |
| **Chr3_BS897_sg2** | CATGTG | 35 | 0 | 0 (cut in the middle) | - |
| **Chr11_BS79_sg1** | CGCGTG | 9 | 40 | 3 | CGCGTGG |
| **Chr11_BS79_sg2** | CGCGTG | 17 | 0 | 0 (cut in the middle) | - |
| **Chr2_BS1664_sg1** | CACGTG | 30 | 70 | 28 | CACGTGG |
| **Chr19_BS2255_sg1** | CACATG | 35 | 0 | 0 (cut in the middle) | - |
| **Chr19_BS2255_sg2** | CACATG | 60 | 0 | 0 (cut in the middle) | - |
| **Chr13_BS121_sg1** | CGCGTG | 0 | 0 | 0 | - |
| **Chr13_BS121_sg2** | CGCGTG | 55 | 62 | 34 | CCGCGTG |

**Table S9.** Mutations of K562 clones.

| Name | Mutations |
|---|---|
| **B4** | wild type homozygous |
| **F10** | wild type homozygous |
| **C10** | wild type homozygous |
| **D6A** | wild type homozygous |
| **D6B** | wild type homozygous |
| **F6** | wild type homozygous |
| **G9** | wild type homozygous |
| **B2** | homozygous +G |
| **G4A** | homozygous +G |
| **G6A** | homozygous +G |
| **F9** | 2 wild type alleles, deletion -17 ntt |
| **F11** | 2 wild type alleles, deletion -39 nt |
| **D11** | 1 wild type allele, deletions -37 nt and -54 nt |
| **B3** | 1 wild type allele, 2x deletion -49 nt |
| **G3** | 1 wild type allele, 2x deletion -29 nt |
| **G4B** | 2x deletion -17 nt and insertion +G |
| **G7** | 2x deletion -18 nt and insertion +G |
| **C8A** | deletions -15 nt and 2x -31 nt |
| **D2** | deletions -20 nt -19 nt and -4 nt |
| **D7** | Deletions -18 nt and 2x -15 nt |
| **D8** | 2x deletion -41nt and +1 insertion |
| **B9** | deletions -43 nt and 2x -32 nt |
| **C9** | 3x deletion >-200 nt |
| **G8** | 3x deletion >-200 nt |

**Table S10.** Oligo sequences.

| Oligo name | Sequence |
|---|---|
| **sgRNA** | |
| **chr17_BS377_sg1-S** | CACCGACGCGGTAACTATACTCACG |
| **chr17_BS377_sg1-AS** | AAACCGTGAGTATAGTTACCGCGTC |
| **chr17_BS377_sg2-S** | CACCGGGAGGATCCAGGTCCGCACG |
| **chr17_BS377_sg2-AS** | AAACCGTGCGGACCTGGATCCTCCC |
| **chr17_BS377_sg3-S** | CACCGGTGAGTATAGTTACCGCGTG |
| **chr17_BS377_sg3-AS** | AAACCACGCGGTAACTATACTCACC |
| **chr11_BS2113_sg1-S** | CACCGAAGCCAGCAGGTGAGACATG |
| **chr11_BS2113_sg1-AS** | AAACCATGTCTCACCTGCTGGCTTC |
| **chr10_BS212_sg1-S** | CACCGTTGGGCGAGAGGGAGACATG |
| **chr10_BS212_sg1-AS** | AAACCATGTCTCCCTCTCGCCCAAC |
| **chr11_BS79_sg1-S** | CACCGCGGTCACCACACTCACGCGG |
| **chr11_BS79_sg1-AS** | AAACCCGCGTGAGTGTGGTGACCGC |
| **chr11_BS79_sg2-S** | CACCGGCCCGGTCACCACACTCACG |
| **chr11_BS79_sg2-AS** | AAACCGTGAGTGTGGTGACCGGGCC |
| **chr2_BS1664_sg1-S** | CACCGCAAGTAGTAGGCTCGGCACG |
| **chr2_BS1664_sg1-AS** | AAACCGTGCCGAGCCTACTACTTGC |
| **chr19_BS2255_sg1-S** | CACCGAGCGTAGTGACCATCATGTG |
| **chr19_BS2255_sg1-AS** | AAACCACATGATGGTCACTACGCTC |
| **chr19_BS2255_sg2-S** | CACCGCCTAGCCCGGCCTCACATGA |
| **chr19_BS2255_sg2-AS** | AAACTCATGTGAGGCCGGGCTAGGC |
| **chr13_BS121_sg1-S** | CACCGAGCGCCCGGGAGCCACGCGT |
| **chr13_BS121_sg1-AS** | AAACACGCGTGGCTCCCGGGCGCTC |
| **chr13_BS121_sg2-S** | CACCGTTCCAGGTAGGGCCTACGCG |
| **chr13_BS121_sg2-AS** | AAACCGCGTAGGCCCTACCTGGAAC |
| **chr3_BS897_sg1-S** | CACCGCGCAGCCGTGGCTCATGTGA |
| **chr3_BS897_sg1-AS** | AAACTCACATGAGCCACGGCTGCGC |
| **chr3_BS897_sg2-S** | CACCGTCGCAGCCGTGGCTCATGTG |
| **chr3_BS897_sg2-AS** | AAACCACATGAGCCACGGCTGCGAC |
| **chr11_BS79_sg3-S** | CACCGCCGCCGGGCGAGGCCGCGCG |
| **chr11_BS79_sg3-AS** | AAACCGCGCGGCCTCGCCCGGCGGC |
| **PFAS_sg1-S** | CACCGAAGCCCATCATGTTTAGTGG |
| **PFAS_sg1-AS** | AAACCCACTAAACATGATGGGCTTC |
| **PFAS_sg2-S** | CACCGTGGACCCAAAAGTCGCCGCC |
| **PFAS_sg2-AS** | AAACGGCGGCGACTTTTGGGTCCAC |
| **PRKCQ_sg1-S** | CACCGCGATGATGTTGAGTGCACGA |
| **PRKCQ_sg1-AS** | AAACTCGTGCACTCAACATCATCGC |
| **PRKCQ_sg2-S** | CACCGGCTCCATCAAAAATGAAGCA |
| **PRKCQ_sg2-AS** | AAACTGCTTCATTTTTGATGGAGCC |
| **RPLP2_sg1-S** | CACCGGGACAGCGTGGGTATCGAGG |
| **RPLP2_sg1-AS** | AAACCCTCGATACCCACGCTGTCCC |
| **RPLP2_sg2-S** | CACCGGGACGACGACCGGCTCAACA |
| **RPLP2_sg2-AS** | AAACTGTTGAGCCGGTCGTCGTCCC |
| **PIDD1_sg1-S** | CACCGAGGGCGTCATGAGGACCCAG |
| **PIDD1_sg1-AS** | AAACCTGGGTCCTCATGACGCCCTC |
| **PIDD1_sg2-S** | CACCGGGGGGCGTCTAGCAGCTCAG |
| **PIDD1_sg2-AS** | AAACCTGAGCTGCTAGACGCCCCCC |
| **shRNA** | |
| **NT2-S** | GATCCGCAACAAGATGAAGAGCACCAACTCTTCAAGAGAGTTGTTCTACTTCTCGTGGTTGAGTTTTTG |
| **NT2-AS** | GCGTTGTTCTACTTCTCGTGGTTGAGAAGTTCTCTCAACAAGATGAAGAGCACCAACTCAAAAACTTAA |
| **PRKCQ-AS1_sh1-S** | GATCCACGCTAGAAAGGGCTTGTAAATTCAAGAGATTTACAAGCCCTTTCTAGCGTTTTTTG |
| **PRKCQ-AS1_sh1-AS** | AATTCAAAAAACGCTAGAAAGGGCTTGTAAATCTCTTGAATTTACAAGCCCTTTCTAGCGTG |
| **PRKCQ-AS1_sh2-S** | GATCCGGGATTTAGGATAGAAATTAATTCAAGAGATTAATTTCTATCCTAAATCCCTTTTTG |
| **PRKCQ-AS1_sh2-AS** | AATTCAAAAAGGGATTTAGGATAGAAATTAATCTCTTGAATTAATTTCTATCCTAAATCCCG |
| **MYC_sh1-S** | GATCCGATGAGGAAGAAATCGATGTTCAAGAGACATCGATTTCTTCCTCATCTTTTTG |
| **MYC_sh1-AS** | AATTCAAAAAGATGAGGAAGAAATCGATGTCTCTTGAACATCGATTTCTTCCTCATCG |
| **MYC_sh3-S** | GATCCAACGACGAGAACAGTTGAAACATTCAAGAGATGTTTCAACTGTTCTCGTCGTTTTTTTG |
| **MYC_sh3-AS** | AATTCAAAAAAACGACGAGAACAGTTGAAACATCTCTTGAATGTTTCAACTGTTCTCGTCGTTG |

| Oligo name | Sequence |
|---|---|
| **MYC responsive element** | |
| **MYC_RE_S** | CCACGTGCACGTGCACGTGCACGTGCACGTGCACGTGC |
| **MYC_RE_AS** | TCGAGCACGTGCACGTGCACGTGCACGTGCACGTGCACGTGGAGCT |

**Table S11.** Primer sequences.

| Primer name | Sequence |
|---|---|
| **sgRNA LIBRARY AMPLIFICATION** | |
| **oligo-F** | GTAACTTGAAAGTATTTCGATTTCTTGGCTTTATATATCTTGTGGAAAGGACGAAACACC |
| **oligo-R** | ACTTTTTCAAGTTGATAACGGACTAGCCTTATTTTAACTTGCTATTTCTAGCTCTAAAAC |
| **NGS LIBRARY PREPARATION** | |
| **Fwd-1** | AATGATACGGCGACCACCGAGATCTACACTCTTTCCCTACACGACGCTCTTCCGATCTTAAGTAGAGGCTTTATATATCTTGTGGAAAGGACGAAACACC |
| **Fwd-2** | AATGATACGGCGACCACCGAGATCTACACTCTTTCCCTACACGACGCTCTTCCGATCTATCATGCTTAGCTTTATATATCTTGTGGAAAGGACGAAACACC |
| **Fwd-3** | AATGATACGGCGACCACCGAGATCTACACTCTTTCCCTACACGACGCTCTTCCGATCTGATGCACATCTGCTTTATATATCTTGTGGAAAGGACGAAACACC |
| **Fwd-4** | AATGATACGGCGACCACCGAGATCTACACTCTTTCCCTACACGACGCTCTTCCGATCTCGATTGCTCGACGCTTTATATATCTTGTGGAAAGGACGAAACACC |
| **Fwd-5** | AATGATACGGCGACCACCGAGATCTACACTCTTTCCCTACACGACGCTCTTCCGATCTTCGATAGCAATTCGCTTTATATATCTTGTGGAAAGGACGAAACACC |
| **Fwd-6** | AATGATACGGCGACCACCGAGATCTACACTCTTTCCCTACACGACGCTCTTCCGATCTATCGATAGTTGCTTGCTTTATATATCTTGTGGAAAGGACGAAACACC |
| **Fwd-7** | AATGATACGGCGACCACCGAGATCTACACTCTTTCCCTACACGACGCTCTTCCGATCTGATCGATCCAGTTAGGCTTTATATATCTTGTGGAAAGGACGAAACACC |
| **Fwd-8** | AATGATACGGCGACCACCGAGATCTACACTCTTTCCCTACACGACGCTCTTCCGATCTCGATCGATTTGAGCCTGCTTTATATATCTTGTGGAAAGGACGAAACACC |
| **Fwd-9** | AATGATACGGCGACCACCGAGATCTACACTCTTTCCCTACACGACGCTCTTCCGATCTACGATCGATACACGATCGCTTTATATATCTTGTGGAAAGGACGAAACACC |
| **Fwd-10** | AATGATACGGCGACCACCGAGATCTACACTCTTTCCCTACACGACGCTCTTCCGATCTTACGATCGATGGTCCAGAGCTTTATATATCTTGTGGAAAGGACGAAACACC |
| **Rev-1** | CAAGCAGAAGACGGCATACGAGATTCGCCTTGGTGACTGGAGTTCAGACGTGTGCTCTTCCGATCTCCGACTCGGTGCCACTTTTTCAA |
| **Rev-2** | CAAGCAGAAGACGGCATACGAGATATAGCGTCGTGACTGGAGTTCAGACGTGTGCTCTTCCGATCTCCGACTCGGTGCCACTTTTTCAA |
| **Rev-3** | CAAGCAGAAGACGGCATACGAGATGAAGAAGTGTGACTGGAGTTCAGACGTGTGCTCTTCCGATCTCCGACTCGGTGCCACTTTTTCAA |
| **Rev-4** | CAAGCAGAAGACGGCATACGAGATATTCTAGGGTGACTGGAGTTCAGACGTGTGCTCTTCCGATCTCCGACTCGGTGCCACTTTTTCAA |
| **Rev-5** | CAAGCAGAAGACGGCATACGAGATCGTTACCAGTGACTGGAGTTCAGACGTGTGCTCTTCCGATCTCCGACTCGGTGCCACTTTTTCAA |
| **Rev-6** | CAAGCAGAAGACGGCATACGAGATGTCTGATGGTGACTGGAGTTCAGACGTGTGCTCTTCCGATCTCCGACTCGGTGCCACTTTTTCAA |
| **Rev-7** | CAAGCAGAAGACGGCATACGAGATTTACGCACGTGACTGGAGTTCAGACGTGTGCTCTTCCGATCTCCGACTCGGTGCCACTTTTTCAA |
| **Rev-8** | CAAGCAGAAGACGGCATACGAGATTTGAATAGGTGACTGGAGTTCAGACGTGTGCTCTTCCGATCTCCGACTCGGTGCCACTTTTTCAA |
| **TIDE PRIMERS** | |
| **chr17_BS377_TIDE-F** | CCCTGATCTTGCCAAGCAGA |
| **chr17_BS377_TIDE-R** | CCCTCCCCAACTTTCAGGAC |
| **chr10_BS212_TIDE-F** | ACCCGGCACCTCTAGCCA |
| **chr10_BS212_TIDE-R** | CCCGCAAGGCATAAAAAGTCC |
| **chr11_BS79_TIDE-F** | AGTCACTTCCGGAACTGCTG |
| **chr11_BS79_TIDE-R** | CCCACGCTGTCCAAGATCTT |
| **chr19_BS2255_TIDE-F** | CCTCTCCCGAAGTCACGATG |
| **chr19_BS2255_TIDE-R** | CGGCGGGCATTTGGAAATAG |
| **chr13_BS121_TIDE-F** | CGGTTGCCTTTCTTTCGCAA |
| **chr13_BS121_TIDE-R** | AGCCGACGCCTGACTTTAAA |
| **chr11_BS2113_TIDE-F** | GCACACCGCTTTGTCATTGT |
| **chr11_BS2113_TIDE-R** | TCAGTTCCGCTTGAGGCATT |
| **chr2_BS1664_TIDE-F** | CCCTGGCTTCAGCAGAATCA |
| **chr2_BS1664_TIDE-R** | GGACACAGGACCAGCTTCTG |
| **chr3_BS897_TIDE-F** | AGGAAGAGCTTTCTGGTGGC |

| Primer name | Sequence |
| --- | --- |
| **chr3_BS897_TIDE-R** | GCTTGGACCCTTTCTCCTCC |
| **qPCR PRIMERS** | |
| **RPLP2-F** | GATCTTGGACAGCGTGGGTA |
| **RPLP2-R** | GCAATGACGTCTTCAATGTTTTT |
| **SNRPD2-F** | GAAACGGGAGTGAACGGAG |
| **SNRPD2-R** | TGGGGTCATCTCACTCTTGG |
| **QPCTL-F** | AACTGGATCCACAGCGTCTC |
| **QPCTL-R** | AAGAGCAGTTGCAGGGTCAC |
| **CTC1-F** | CAGCTGTCACCCACGTGTC |
| **CTC1-R** | TGGACACTCGCAGTTCTGTC |
| **PRKRIR-F** | CTTCCTGCCTTATGAAGCCGA |
| **PRKRIR-R** | CCTGGCCACGACAATACTCC |
| **PRKCQ-F** | CAAGTGCCACGAGTTCACTG |
| **PRKCQ-R** | GCACTGGTAGCCCTGTTTGT |
| **PFAS-F** | GTGAGTGGATCAAGCCCATC |
| **PFAS-R** | GTAGACGGGACCTCCAACCT |
| **PIDD1-F** | CTAGACGCCCCCTTTGTGC |
| **PIDD1-R** | GGTCAGGAACAGTCTGGGC |
| **PRKCQ-AS1-F** | CCCACAACTCGGAACTTGGA |
| **PRKCQ-AS1-R** | AGGAAGGATGCAAGACGTGG |
| **LINC00412-F** | CGGGGTTGTCACTGGAAGTT |
| **LINC00412-R** | CAGCAAAGTTGTACCATGACCC |
| **RPL21-F** | CCTTTGGCCACATATATGCGAATC |
| **RPL21-R** | ACACTTGTGGGGCATTCCTT |
| **SCAP-F** | ATCTCGGGCCTTCTACAACC |
| **SCAP-R** | CAAGGGGAGTTTCAGCAGTG |
| **PTPN23-F** | GAGGGCATGAAGGTCTCCTG |
| **PTPN23-R** | GCATGAGGTTGACGTTGAGC |
| **TBP-F** | GCCCGAAACGCCGAATAT |
| **TBP-R** | CCGTGGTTCGTGGCTCTCT |
| **LUCIFERASE CONSTRUCTS** | |
| **chr10_BS212_luc-F** | ATATGTGAGCTCCGCTCGCCAGCCTCC |
| **chr10_BS212_luc-R** | ATATGTCTCGAGCAGATACAGGAAAAGTCCCAGGT |
| **chr11_BS79_luc-F** | TAACGAGAGCTCGGTGCTAGGTACCGAAGGC |
| **chr11_BS79_luc-R** | TAACGACTCGAGAAAAATACCCCGCCGCCC |
| **chr17_BS377_luc-F** | ATTCGTGAGCTCTGCTTGCTTTATGGCCTTAACTAAC |
| **chr17_BS377_luc-R** | ATTCGTCTCGAGTGAGGGCTGTATTCCATGACC |
| **ChIP-qPCR** | |
| **chr10_BS212_ChIP-F** | CCTCTCGCCCAACTGAAAAC |
| **chr10_BS212_ChIP-R** | CAGAGAAAAGAGACCCAGCTC |
| **chr11_BS79_ChIP-F** | GTCCCTTTGGACTCGCTTC |
| **chr11_BS79_ChIP-R** | GTTCCGGAAGTGACTGCTCT |
| **chr17_BS377_ChIP-F** | TTTCGTCATCTAGCCCAAGC |
| **chr17_BS377_ChIP-R** | GGGCACCAAGTAGACACAGC |

**Table S12.** Number of reads obtained by NGS for individual samples.

| NGS pool | Sample | # reads | # perfect sgRNA matches | Coverage |
|---|---|---|---|---|
| **1** | MYC-CRISPR library plasmid | 116 452 060 | 106 981 131 | 2 308x |
| | Brunello library plasmid | 157 895 429 | 144 707 272 | 1 868x |
| **2** | K562_MYC-CRISPR #1 T0 | 83 489 199 | 75 443 224 | 1 627x |
| | K562_MYC-CRISPR #1 T1 | 59 895 534 | 53 260 435 | 1 149x |
| | K562_MYC-CRISPR #2 T0 | 63 067 319 | 56 420 523 | 1 217x |
| | K562_MYC-CRISPR #2 T1 | 58 484 870 | 52 205 808 | 1 126x |
| **3** | K562_Brunello #1 T0 | 54 221 760 | 49 649 571 | 641x |
| | K562_Brunello #1 T1 | 65 892 957 | 58 862 286 | 760x |
| | K562_Brunello #2 T0 | 72 332 476 | 66 210 988 | 855x |
| | K562_Brunello #2 T1 | 66 241 094 | 59 146 212 | 764x |
| **4** | ST486_MYC-CRISPR #1 T0 | 79 321 637 | 72 156 055 | 1 556x |
| | ST486_MYC-CRISPR #1 T1 | 87 870 982 | 79 467 416 | 1 714x |
| | ST486_MYC-CRISPR #2 T0 | 59 934 111 | 54 499 953 | 1 175x |
| | ST486_MYC-CRISPR #2 T1 | 70 122 820 | 63 352 176 | 1 366x |
| **5** | ST486_Brunello #1 T0 | 70 116 091 | 63 554 293 | 820x |
| | ST486_Brunello #1 T1 | 89 895 834 | 80,185,600 | 1 035x |
| | ST486_Brunello #2 T0 | 70,283,191 | 63 729 565 | 823x |
| | ST486_Brunello #2 T1 | 75 121 180 | 67 041 436 | 865x |
| **6** | HepG2_MYC-CRISPR #1 T0 | 51 447 782 | 47 155 897 | 1 017x |
| | HepG2_MYC-CRISPR #1 T1 | 66 626 438 | 60 555 248 | 1 306x |
| | HepG2_MYC-CRISPR #2 T0 | 50 858 585 | 46 650 129 | 1 006x |
| | HepG2_MYC-CRISPR #2 T1 | 52 090 164 | 47 337 827 | 1 021x |
| **7** | HepG2_Brunello #1 T0 | 44 194 540 | 33 638 192 | 434x |
| | HepG2_Brunello #1 T1 | 54 746 834 | 49 087 636 | 634x |
| | HepG2_Brunello #2 T0 | 60 630 324 | 55 661 488 | 719x |
| | HepG2_Brunello #2 T1 | 52 464 546 | 47 052 280 | 608x |
| **8** | MCF7_MYC-CRISPR #1 T0 | 55 595 928 | 50 892 310 | 1 098x |
| | MCF7_MYC-CRISPR #1 T1 | 63 075 305 | 57 089 710 | 1 232x |
| | MCF7_MYC-CRISPR #2 T0 | 50 858 585 | 46 650 129 | 1 006x |
| | MCF7_MYC-CRISPR #2 T1 | 50 339 659 | 45 688 415 | 986x |
| **9** | MCF7_Brunello #1 T0 | 53 498 499 | 48 765 805 | 630x |
| | MCF7_Brunello #1 T1 | 68 312 092 | 61 109 079 | 789x |
| | MCF7_Brunello #2 T0 | 55 841 233 | 51 347 850 | 663x |
| | MCF7_Brunello #2 T1 | 45 446 430 | 37 428 965 | 483x |

#### SUPPLEMENTARY FIGURES

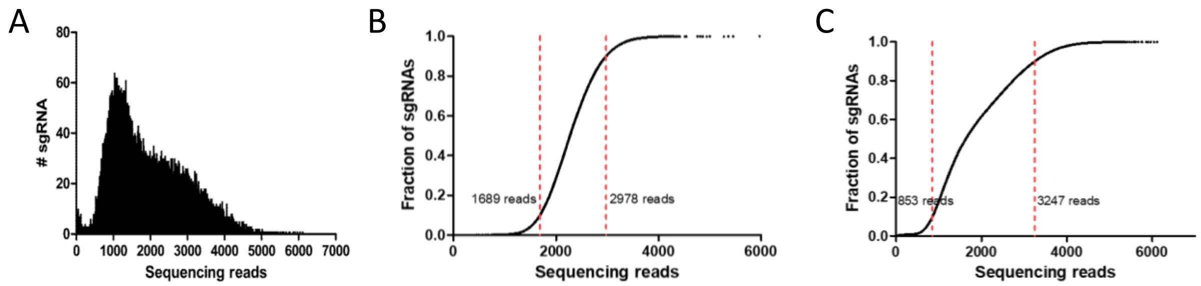


**Figure S1. Quality of MYC-CRISPR and Brunello libraries based on NGS. A)** Distribution of sgRNA read counts in the libraries. **B) and C)** Cumulative frequency of sgRNAs. Red lines indicate the $10^{th}$ and $90^{th}$ percentile in **B)** MYC-CRISPR and **C)** Brunello library.

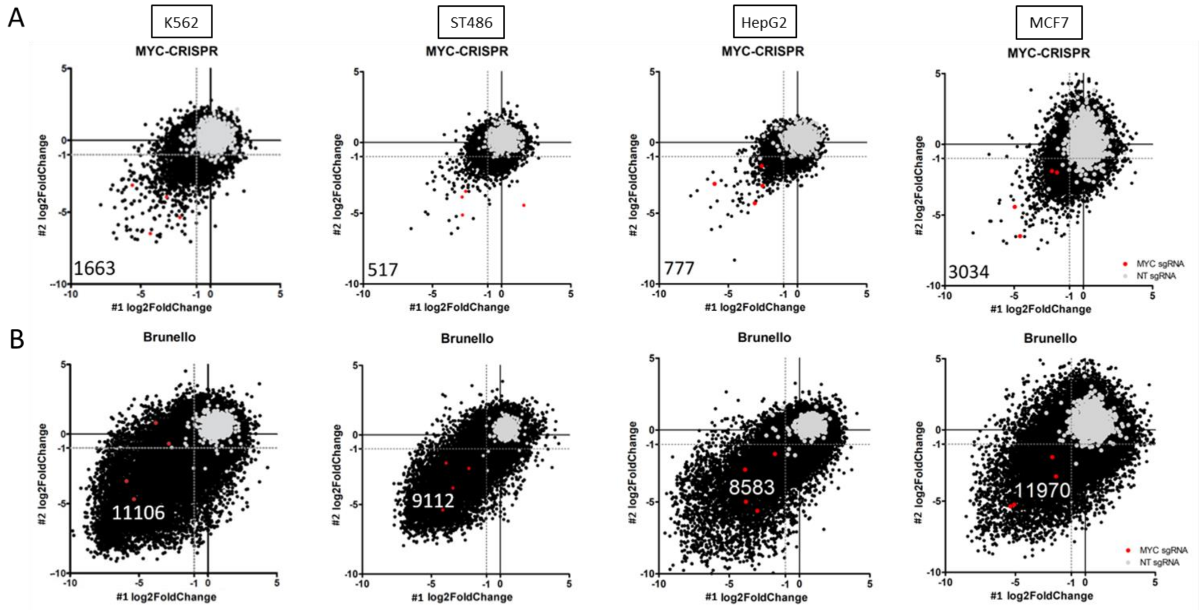


**Figure S2. Changes in sgRNA abundance in two screen replicates with A)** MYC-CRISPR and **B)** Brunello library. Log2 fold change values for replicate #1 and #2 are shown on the X and Y axis.

A

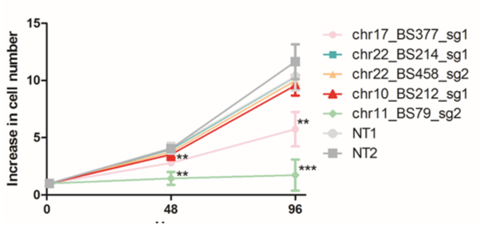


B

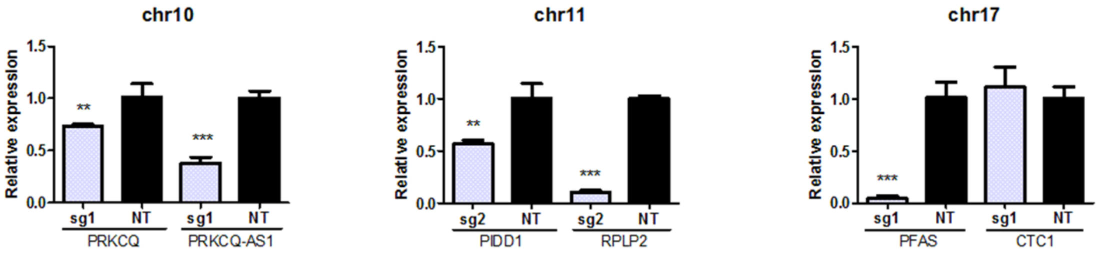


**Figure S3. Validation of the screen results using the CRISPR/dCas9 approach. A)** Cell viability after CRISPR/dCas9 blocking of selected E-boxes. Cells expressing dCas9 were infected with individual sgRNAs targeting E-boxes. After puromycin selection for four days, cell viability was measured using CellTiter-Glo assay at three timepoints: 0, 48 and 96 hours. Shown are average values and standard deviations from three independent experiments, each performed in triplicate. **, p<0.01; ***, p<0.001, Student's t-test **B)** qRT-PCR analysis of expression of genes adjacent to selected E-boxes upon CRISPR/dCas9 blocking of E-box sequences. For all E-boxes at least one nearby gene showed significantly decreased expression. The mean and SD of two independent experiments, each performed in triplicate, are shown. **, p<0.01; ***, p<0.001, Student's t-test.

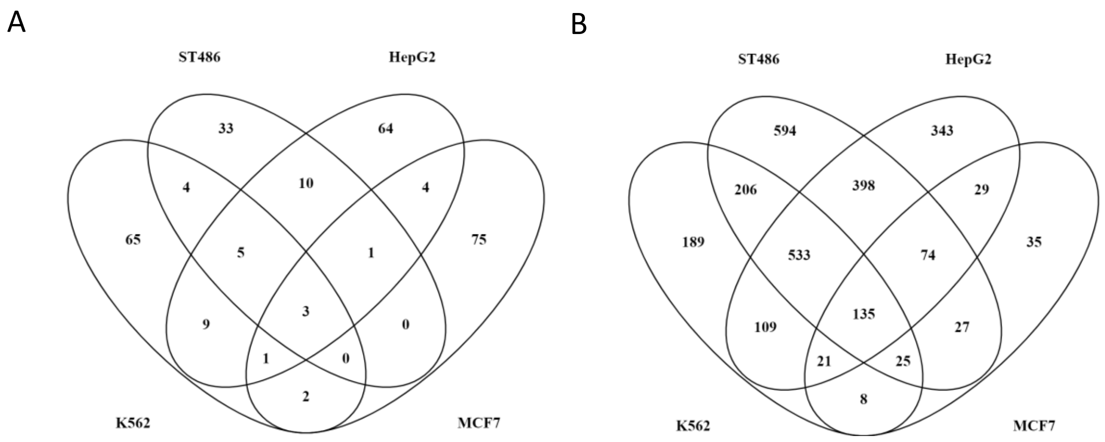


**Figure S4. Overlap of essential A)** E-boxes from MYC-CRISPR and **B)** genes from Brunello library.

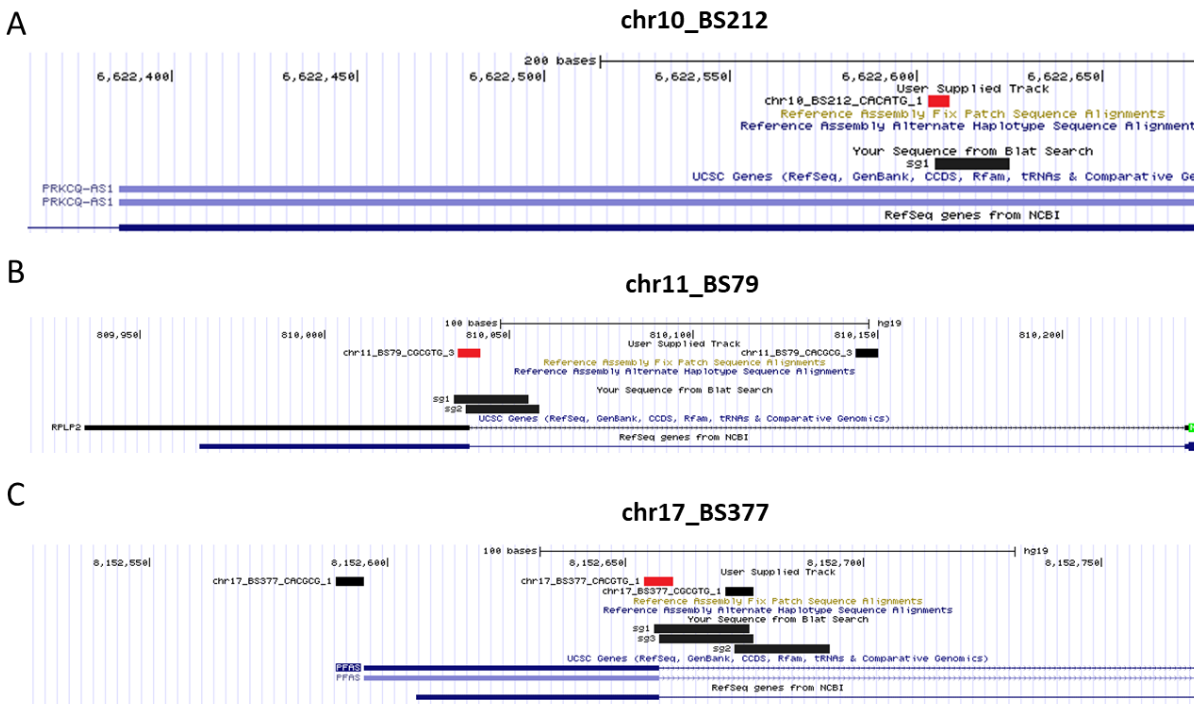


**Figure S5. Genomic location of sgRNAs targeting selected E-boxes based on UCSC hg19.** Essential E-boxes marked in red.

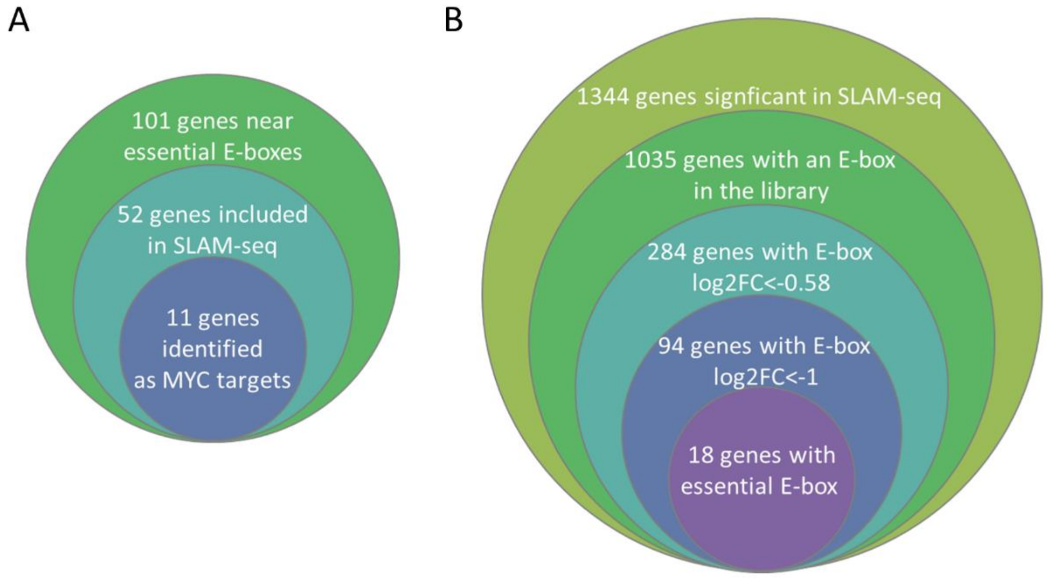


**Figure S6**. **Intersection of the MYC-CRISPR screen with SLAM-seq data**[17]**. A)** From the viewpoint of 101 genes near essential E-boxes identified in the MYC-CRISPR screen. **B)** From the viewpoint of 1344 genes identified as MYC targets in SLAM-seq.
